# Brain-wide synaptosome profiling reveals localized mRNAs that diversify synapses

**DOI:** 10.64898/2025.12.07.692806

**Authors:** Marcel Juengling, Janus Mosbacher, Julio Perez, Marc van Oostrum, Eva Kaulich, Susanne tom Dieck, Nicole Fuerst, Georgi Tushev, Maria Korelidou, Erin M. Schuman

## Abstract

Chemical synapses are the principal communication nodes of the brain, defined by their specialization and plasticity. Their diversity spans multiple levels, from the polarity and magnitude of electrophysiological responses to the proteins driving these differences. Here, we describe another layer of synapse diversity – synaptic transcriptome diversity. Using transgenic mice to fluorescently label presynapses in Camk2a-, Gad2-, DAT-, PV-, SST-, and VIP-expressing neurons, combined with fluorescence-activated synaptosome sorting, we profiled synaptic transcriptomes across five brain regions. We identified ∼4000 mRNAs enriched at synapses – some type-specific, others shared – and highlight region-specific enrichments of mRNAs encoding protein subunits or family members. We found that the abundance of synaptic mRNAs is not a passive reflection of their abundance in cellular somata, indicating active trafficking and sorting mechanisms. Integrating transcriptomic and proteomic data, we identified ∼90 genes with significantly correlated mRNA-protein ratios across regions, mainly involved in synaptic vesicle dynamics, receptor signaling, and calcium regulation, suggesting a key role for local translation in maintaining protein copy number. Although synaptic mRNAs represent only a fraction of the templates for the synaptic proteome, the synaptic transcriptome reflects the synapse diversity captured by the proteome equally well. Together, this dataset (https://syndive.org/) provides a resource for exploring how synaptic mRNA localization and local translation shape synaptic identity and function.

## INTRODUCTION

The elaborate morphology of the neuron, with its vast dendritic and axonal arbor, poses unique challenges for proteostasis and proteome remodeling. This challenge is especially apparent at synapses, which operate far from the nucleus, the site of transcription. The dynamic demands of synaptic function require a diverse array of thousands of proteins (Oostrum et al., 2023), each with varying lifespans and copy numbers (Bulovaite et al., 2022; Dörrbaum et al., 2018; Fornasiero et al., 2018; Wilhelm et al., 2014). The transport of mRNAs to neuronal processes and their subsequent local translation into proteins near or at synapses is a vital mechanism neurons use to meet these demands (Bourke et al., 2023; Chekulaeva, 2024; J. D. Perez, Fusco, et al., 2021). This process ensures a finely tuned temporal and spatial control of the synaptic proteome, which is crucial for synaptic maintenance and plasticity.

Many RNA sequencing studies have characterized the presence and differential abundance of thousands of mRNAs in neurites (axons and dendrites) compared to their somata (Cajigas et al., 2012; Glock et al., 2021; Kaulich et al., 2025; Taliaferro et al., 2016; Zappulo et al., 2017; Zivraj et al., 2010). Ribosome profiling experiments have even demonstrated that for ∼800 proteins, the dominant source of their translation is the neuropil (Glock et al., 2021). These studies and others have highlighted the specialized mechanisms neurons employ for localizing and translating mRNAs, including different isoform usage for localization and stabilization of mRNAs (Taliaferro et al., 2016; Tushev et al., 2018), and monosome translation (Biever et al., 2020). However, it remains unknown to what extent the local transcriptome previously characterized in axons and dendrites reflects the synaptic transcriptome. This distinction is crucial as different mechanisms might influence the localization and function of mRNAs that are *adjacent to* (e.g. within the main axon or dendritic shaft) versus *within* a synapse. Understanding these mechanisms could reveal the specificity of local translation, differentiating its role in synthesizing proteins for synaptic neighborhoods vs. individual synapses (Rangaraju et al., 2017; Sun et al., 2021). It could also elucidate whether the mRNAs present at different synapse types encode synaptic proteins that are common to all synapses, or alternatively, encode proteins that specify the uniqueness of a particular synapse type. The population of localized synaptic mRNAs might, further, be a descriptor of functional types and/or the plasticity potential of a given synaptic population **–** both factors which are a function of pre- and postsynaptic cell type, and neural activity. Excitatory and inhibitory synapses, for example, require different molecular machinery to support depolarization versus hyperpolarization or shunting, likely requiring specialized sets of localized mRNAs. Also, more fine-grained neuronal subtypes sharing the same neurotransmitters, like the inhibitory parvalbumin (PV), somatostatin (SST) and vasoactive intestinal peptide (VIP) neurons, may recruit distinct sets of synaptic mRNAs to support their unique firing patterns and connectivity. For example, PV neurons are mainly fast-spiking and target other pyramidal neurons perisomatically; SST neurons are regular-spiking and target pyramidal neuron dendrites. VIP neurons, in turn, are regular-spiking, but primarily target other inhibitory neurons (Tremblay et al., 2016). Characterizing the synaptic transcriptome of different synaptic populations is, therefore, essential for understanding the contribution of local translation to the functional diversification of synapses.

While a few recent publications have highlighted the transcriptomes of purified synaptic populations (Hafner et al., 2019; Hobson et al., 2022; Kaulich et al., 2024; Rubio et al., 2023), a broad and comparative transcriptomic analysis of different synapse types across the mouse brain is still lacking. Here, we use a combination of genetic mouse lines, biochemical fractionation, and Fluorescence-Activated Synaptosome Sorting (FASS) (Biesemann et al., 2014) to isolate purified synapses from Camk2a, several types of Gad2, and DAT neurons to characterize synaptic transcriptomes across the mouse brain.

## RESULTS

### Sequencing synapses of cortical glutamatergic and GABAergic neurons reveals common and distinct transcriptomic signatures

To obtain synapse type-specific transcriptomes, we used our previously established biochemically and electron microscopy-validated pipeline to prepare and purify cell type-specific synaptosomes (isolated synaptic particles containing a resealed presynapse, often attached to a postsynaptic density or resealed postsynapse) from the cerebral cortex (Oostrum et al., 2023). We used Cre-inducible knock-in mice expressing a presynaptic vesicle protein, synaptophysin, fused to tdTomato (SypTOM), to fluorescently label synaptosomes (Figure 1A). Synapses of cortical glutamatergic and GABAergic neurons were labeled in Camk2a-Cre::SypTOM+ (Camk2a) and Gad2-Cre::SypTOM+ (Gad2) mice, respectively. Fluorescence-activated Synaptosome Sorting (FASS) was employed to enrich millions of particles of our synapse types of interest (using the SypTOM and FM dye signal). We also obtained control synaptosomes (using the FM dye signal) originating from the same biochemical preparation but comprising all synapse types and some region-specific membranous contaminants, as previously shown (Figure S1) (Oostrum et al., 2023).

**Figure 1:**
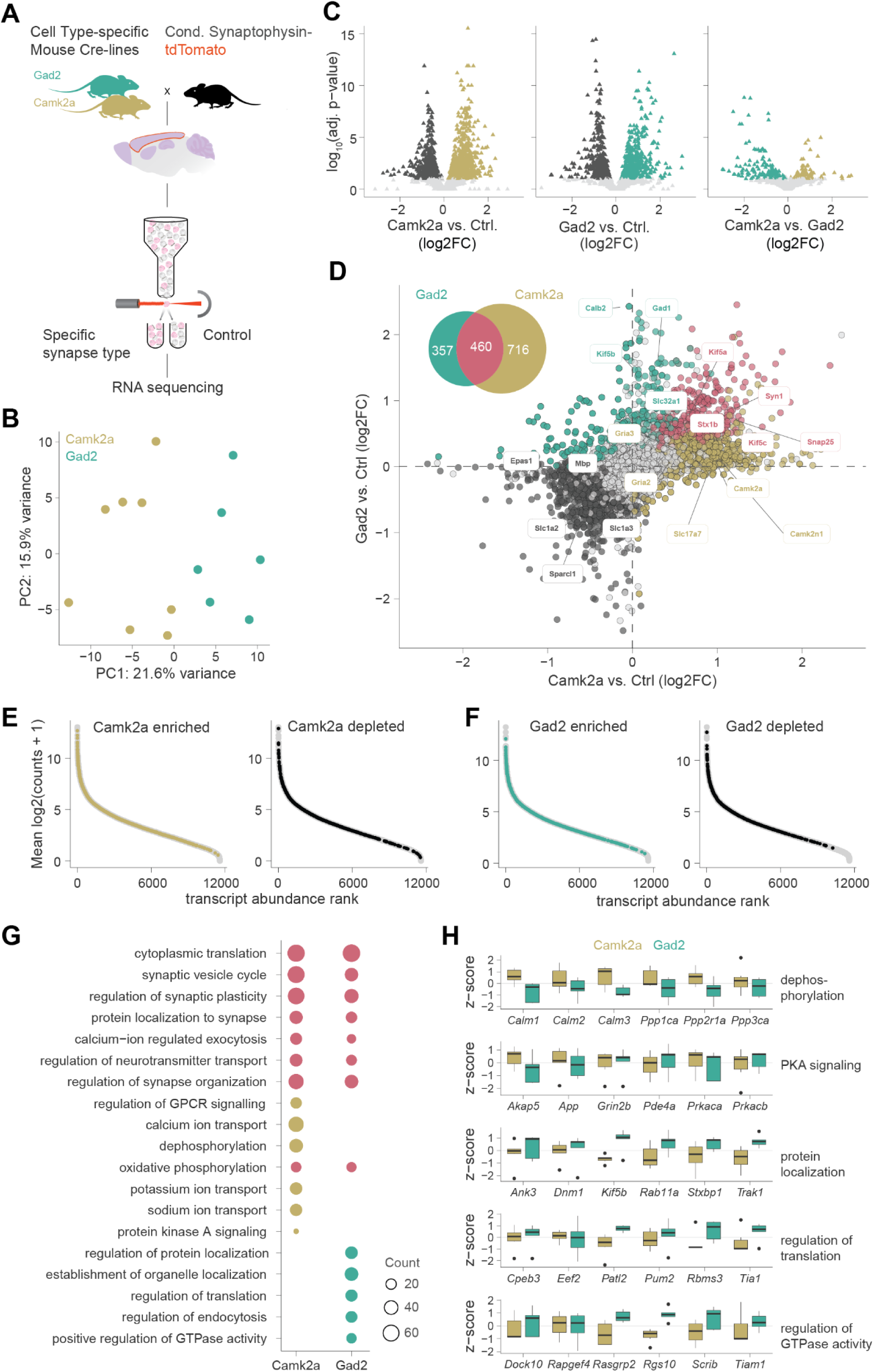
Cell type-specific profiling of synaptic transcriptomes. (A) Workflow schematic adapted from (Oostrum et al., 2023) to isolate cell type-specific synaptosomes. We crossed different Cre-driver lines with a floxed synaptophysin-tdTomato line (SypTOM) and used FASS to enrich our synapse type of interest (tdTomato and FM 4-64 dye signal) and sort a brain region-specific control containing synaptosomes of all types and “synapse-sized” particles (FM dye signal). We then used RNA sequencing to profile synaptic transcriptomes. (B) PCA for the 500 most variable mRNAs of cortical synaptosomes derived from Camk2a and Gad2 animals. Cortical glutamatergic Camk2a and Gad2 synaptosomes exhibit distinct transcriptomic signatures. (C) Volcano plot comparing Camk2a synaptosomes vs. control (left), Gad2 synaptosomes vs. control (middle), and Camk2a vs. Gad2 synaptosomes (right). Significant transcripts are colored in dark grey (control), yellow (Camk2a), and cyan (Gad2). The y-axis is clipped at 15 (D) Scatter plot comparing the differential expression results for Camk2a and Gad2 synaptosomes (|x| and |y| limits < 2.5), highlighting uniquely enriched transcripts (716 for Camk2a, 357 for Gad2), transcripts enriched in both synapse types (460; in pink) and depleted transcripts (dark grey). Synapse type markers like *Camk2a* and *Camk2n1* for Camk2a synaptosomes and *Gad1* and *Slc32a1* for Gad2 synaptosomes are enriched in the respective transcriptomes, whereas, e.g., astrocytic (*Slc1a2*) and nuclear markers (*Epas1*), respectively, are depleted. (E) Normalized mRNA abundance rank for cortical Camk2a synaptosome data. Enriched (left; in yellow) or depleted (right; in black) are indicated. (F) same as (E) for Gad2. (G) Dotplots of selected terms enriched in a GO analysis (category biological process), using the Camk2a and Gad2-enriched mRNAs as input. Commonly shared terms are colored in pink, Camk2a and Gad2 unique terms are marked in yellow and cyan, respectively. (H) Z-scored mRNA levels for selected genes, contributing to the enrichment of uniquely enriched GO terms.

Next, we performed RNA sequencing on 2-10 million sorted cortical synaptosomes and control particles. We used a highly reproducible low-input RNA-sequencing method for synaptosomes (see methods and Table S1 for details; Figures S2, S3A; (Kaulich et al., 2025)), tagging RNAs with a sample index and a UMI, which allows for pooling samples after PCR. We detected 11220 and 11434 types of mRNAs for Camk2a and Gad2 synaptosomes, respectively, after applying an inclusion criterion based on reproducible mRNA detection (see methods). Differential expression analysis revealed only negligible differences attributable to either the sex or strain of the mice (Figure S3B).

Are the excitatory and inhibitory cortical synaptic transcriptomes distinct from one another, or is there a largely common pool of synaptic mRNAs? We observed a clear separation, that is, distinct transcriptomes, for Camk2a and Gad2 synaptosomes using principal component analysis (PCA) (Figure 1B). Comparing Camk2a and Gad2 synaptosomes to their respective “bulk synaptosome” controls and to one another, we detected 100s of differentially expressed mRNAs in each comparison, including many expected synaptic and neuronal markers (Figures 1C,D). Uniquely enriched Camk2a synaptosome mRNAs (∼700) included excitatory markers like the calmodulin-dependent protein kinase *Camk2a*, its respective inhibitor *Camk2n1*, the vesicular glutamate transporter *Slc17a7* and the AMPA receptor subunits *Gria2* and *Gria3*. Conversely, uniquely enriched Gad2 synaptosome mRNAs (∼350) included well-known inhibitory neuron markers like the calcium-binding protein Calretinin (*Calb2*) as well as *Gad1* and *Slc32a1*, which are involved in the synthesis and synaptic vesicle uptake of GABA, respectively. We also observed a population of (∼450) mRNAs enriched in both synapse types, including pan-neuronal mRNAs encoding presynaptic proteins involved in synaptic transmission like *Syn1*, *Snap25*, and *Stx1b*. Importantly, we also observed the enrichment of many mRNAs not typically associated with neuron or synapse identity. For instance, all members of the kinesin-1 heavy chain motor protein family were synaptically enriched, with *Kif5b* showing unique enrichment in Gad2 synaptosomes. This reflects the potential for local translation to supply synapses with proteins involved in plus-end directed transport of protein or organelle cargos. Verifying the enrichment of synaptic compartments in our pipeline, we also detected 100s of depleted mRNAs, including mRNAs for extracellular matrix proteins (e.g., *Sparcl1*) or non-neuronal markers (e.g., the endothelial transcription factor *Epas1*, the astrocytic glutamate transporter *Slc1a2/Slc1a3* or the oligodendrocyte marker *Mbp*) (Figure 1D). Importantly, we found that neither the enrichment nor depletion of a transcript can be predicted by its abundance in a particular (Camk2a or Gad2) synaptosome preparation; mRNAs that were significantly enriched or depleted populated the entire abundance range (Figure 1E,F).

Gene ontology (GO) analysis revealed the pan-synaptic enrichment of terms associated with synapse organization and plasticity, neurotransmitter release, protein transport, mitochondrial function, and protein synthesis (Figure 1G). Excitatory synapses were enriched in mRNAs encoding proteins associated with membrane potential regulation (calcium, sodium, and potassium ion channels), second messenger signaling (G protein-coupled receptors), and phosphorylation (Figure 1G,H). In inhibitory synapses, we observed the enrichment of terms related to localization of proteins and organelles, regulation of translation, endocytosis, and GTPase activity (Fig.1 G,H). These data (Table S2) indicate cortical excitatory and inhibitory synaptic transcriptomes both exhibit common and distinct signatures, suggesting a role for local translation in supplying proteins for general synapse function as well as synapse type specialization.

### Compartment-specific mRNA preferences in synapses compared to somata

How much information do the distinct transcriptomic signatures observed in cortical excitatory and inhibitory synapses carry about their parent cell type, that is, to what extent does the synaptic transcriptome reflect the somatic transcriptome? To answer this, we compared the synaptic transcriptomes to published single-soma (often referred to as “single-cell”) transcriptome data from mouse glutamatergic neurons, GABAergic neurons, and non-neuronal cells (Yao et al., 2021) (Figure 2A, Table S3), and mapped the synaptic mRNAs onto the single-soma maps. In general, we observed that synaptically enriched mRNAs differentiated neuronal from non-neuronal somata, assessed based on average expression across neuron clusters (Figures 2B-D left) and compared with randomly selected mRNAs within a neuron cluster (Figures 2B-D right). As expected, pan-synaptically enriched mRNAs mapped to both glutamatergic and GABAergic neurons (Figure 2B). In contrast, synaptically enriched excitatory and inhibitory mRNAs showed substantially higher levels in their respective neuron types compared to other clusters (Figure 2C,D), showing that the synaptic transcriptome does significantly reflect the parent cell identity. To what extent, though, is the abundance of the individual mRNAs detected at synapses a passive reflection of the somatic mRNA abundance? We plotted the ranked abundance of all overlapping mRNAs in each compartment and compared them. We found that, overall, the somatic abundance of a transcript was a poor predictor of its relative abundance at either excitatory or inhibitory synapses (Figure 2E,F). To quantify this, we calculated the z-score difference between synaptic and somatic expression, with positive values indicating synaptic overrepresentation and negative values indicating somatic overrepresentation of a particular transcript (Figure 2G,H). Interestingly, mRNAs related to synaptic transmission and vesicle dynamics (e.g., *Syn2*, *Rims1*, *Cplx1*) as well as calcium signaling and homeostasis (e.g., *Hpcal4*, *Camk2a*, *Ncald*) showed much higher relative abundance in synapses than in somata. Conversely, several regulators of neuronal excitability and signaling (e.g., the ion channels and transporters *Atp1a1*, *Kcnv1*, *Trpc6*), and GABAergic markers (*Gad1*, *Slc32a1*) showed elevated somatic abundance, likely reflecting need-dependent transcript localization.

**Figure 2:**
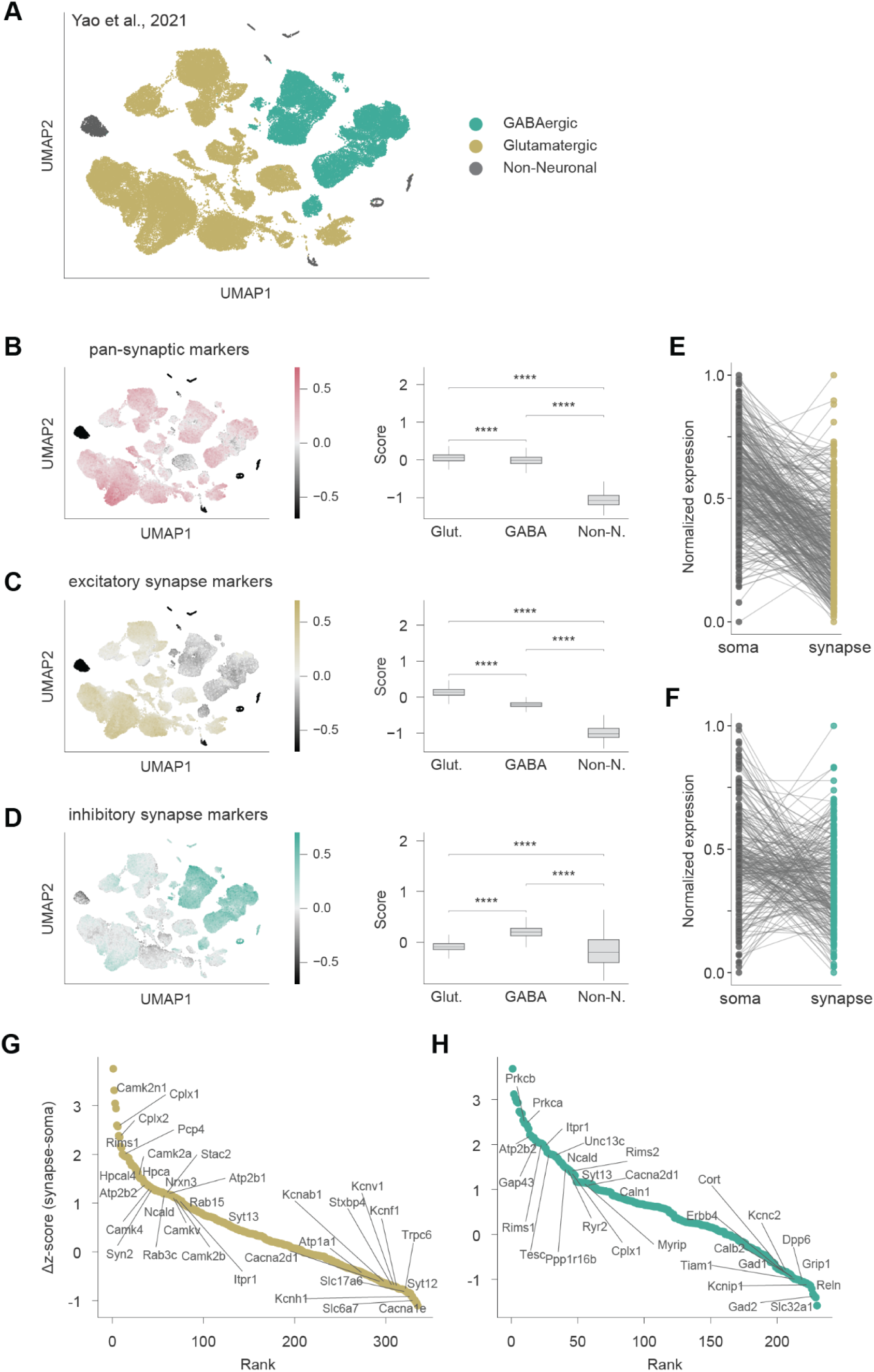
Comparison of the mouse cortical somatic and synaptic transcriptome. (A) UMAP of single-soma sequencing data (Yao et al., 2021) labeled according to cell type. (B) Left: UMAP colored by module scores for pan-synaptically enriched mRNAs (difference between the average expression of pan-synaptic mRNA and an expression-matched control gene set in single-soma data). Right: quantification of module scores in Glutamatergic, GABAergic and Non-Neuronal cells. Dunn’s post-hoc tests (Benjamini-Hochberg-corrected, p-values < 1⋅10^−4^) was used to assess significant differences between categories (C) same as (B), but for uniquely Camk2a-enriched mRNAs. (C) Same as (B) and (C), but for uniquely inhibitory mRNAs. (E) Parallel coordinates plot comparing the mRNA abundance ranks of synaptically enriched genes for glutamatergic soma vs. Camk2a synaptosomes (spearman correlation is 18%). (F) same as (E), but for Gad2. Spearman correlation is −18%. (G) Ranking of the Δz-score of excitatory synapse-enriched transcripts (comparing synaptic and somatic mRNA levels). Selected mRNAs are shown. Positive values denote relative enrichment in synapses compared to somata. (H) Same as G, but for inhibitory synapse-enriched transcripts.

### The 3’UTR landscape of cortical excitatory and inhibitory synapses

While the role of some 3’UTRs in the localization of mRNAs to neurites has been described (An et al., 2008; Andreassi et al., 2021; Grzejda et al., 2025; Mattioli et al., 2019; Tushev et al., 2018), less is known about cell type differences in cis-acting elements that drive the localization of transcripts. We thus compared the 3’UTRs of excitatory and inhibitory synapse-enriched mRNAs. We detected 3’UTRs in our 3’-biased sequencing data using a custom pipeline (increasing the captured 3’UTR diversity compared to current annotations in the NCBI RefSeq database (Goldfarb et al., 2024) (see methods; Figure S4). The mRNAs enriched in Camk2a or Gad2 synaptosomes, or both synapse types, exhibited a large repertoire of 3’UTR isoforms, with around ∼60% of genes expressing more than one mRNA isoform and ∼10% expressing up to four isoforms (Figure 3A). Assessing differential 3’UTR isoform usage, we found that despite this diversity, localized mRNAs enriched in both excitatory and inhibitory synapses used the same 3’UTR isoform (Figure 3B). This suggests that a general need for subcellular localization, rather than neuron type identity, determines the usage of cis-acting localization elements in the 3’UTRs.

**Figure 3:**
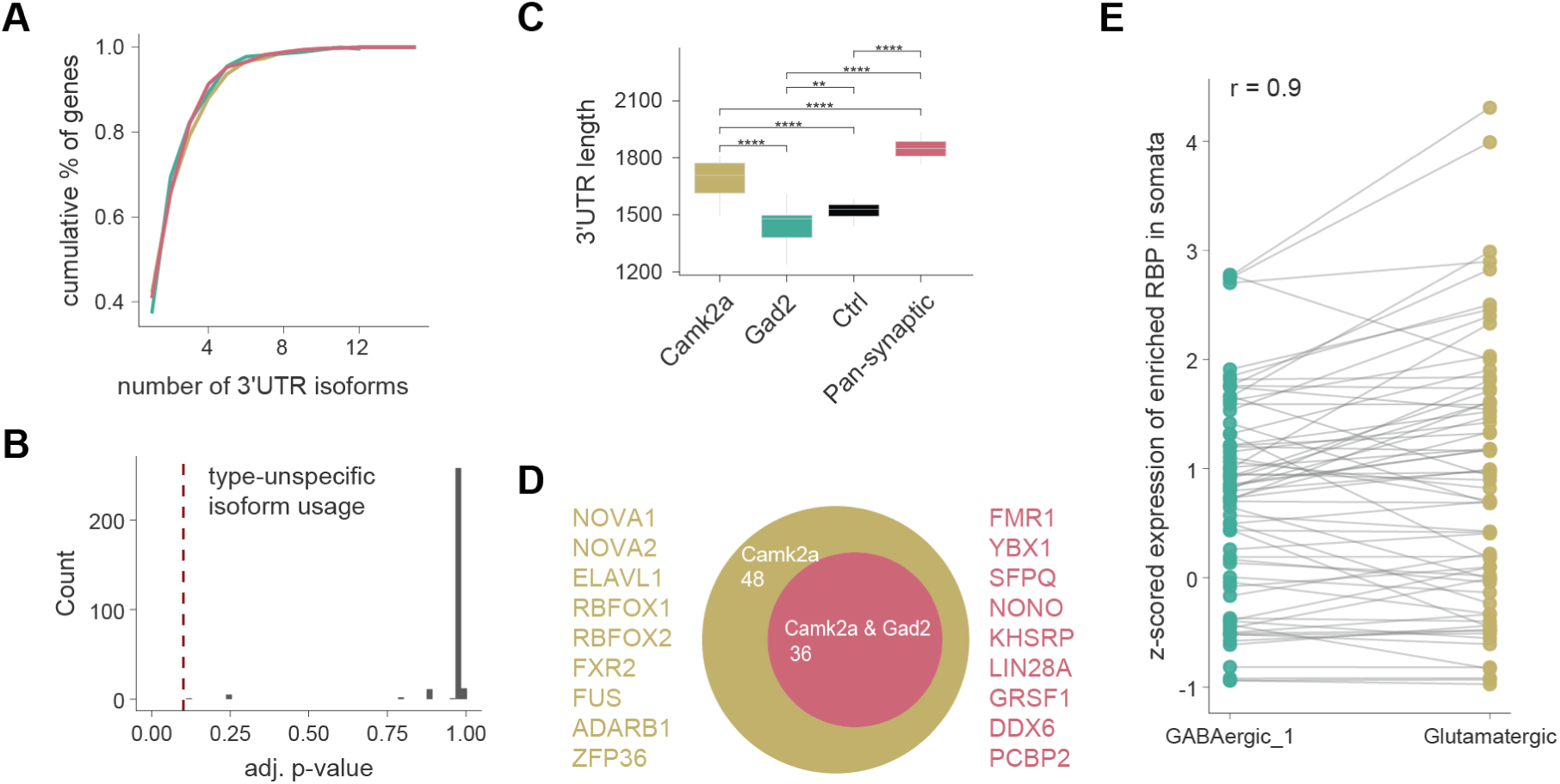
The synaptic 3’UTR regulatory landscape. (A) Cumulative percentage of genes with a given number of 3’UTRs for pan-synaptically (pink), Camk2a (yellow) or Gad2 (cyan) enriched transcripts. (B) Histogram of Benjamini-Hochberg adjusted p-values from the DRIMSeq likelihood ratio test to assess differential isoform usage. The red dashed line shows the significance cutoff (0.1). No type-specific isoform usage was detected. (C) Boxplot showing significant differences between the average 3’ UTR lengths for Camk2a-, Gad2-, and pan-synaptically enriched mRNAs, weighted by expression of the detected 3’UTR isoforms (one-way ANOVA p-value < 2×10⁻6; Tukey’s HSD p-value < 4×10⁻³ for all pairwise comparisons). (D) Venn Diagram of RBPs for which sequence motifs, that is., potential RBP motif binding sites, are enriched in the 3’UTRs of synaptically enriched transcripts. Example RBPs are shown. (E) z-scored expression comparison for RBPs of GABAergic and Glutamatergic somata (J. D. Perez, Dieck, et al., 2021).

One mechanism that affects the potential interaction of mRNA molecules with RNA-binding proteins (RBPs) is the inclusion or exclusion of cis-regulatory elements by lengthening or shortening 3’UTRs (Andreassi et al., 2021; Tushev et al., 2018). Longer 3’UTRs exhibit increased “competitiveness” for regulatory factors by introducing additional motifs for the same or different RBPs (or miRNAs) (Tushev et al., 2018). Comparing the 3’UTRs of synaptically enriched mRNAs, we found that uniquely Camk2a and pan-synaptically enriched mRNAs were longer than those mRNAs that were not synaptically enriched (Figure 3C). Interestingly, the uniquely enriched Gad2 3’UTR isoforms were, on average, shorter than both the other synaptic and control mRNAs. To assess how these length differences correlate with potential RBP interactions, we calculated which RBP motifs are enriched in the 3′UTRs of inhibitory and excitatory synapses. We found substantial overlap in the potential for RBP binding, but also observed that the longer 3’UTRs of excitatory mRNAs can serve as binding platforms for additional RBPs (Figure 3D). Several of the shared and unique motifs were associated with RBPs involved in regulating synaptic function. Synapse type-specificity for RBPs is largely not established. However, we found the motifs for the RNA editing enzyme ADARB1, which has been shown to play a role in functionalizing GluA2 AMPA receptor subunits (Higuchi et al., 2000), and FXR2, which stabilizes GluA1 mRNA (Guo et al., 2015), to be enriched in excitatory 3’UTRs. Generally, we observed motifs for RBPs involved in most facets of posttranscriptional regulation, ranging from RNA stability (e.g., ELAVL1, ZFP36) to RNA transport (e.g., SFPQ), translational regulation (e.g., EIF3D) and the regulation of RNA modifications such m6A methylation (RBM15) or demethylation (FTO), highlighting the potential for local control of RNA translation and metabolism. We also found motifs enriched for two key RBPs associated with synaptopathies – the translational repressor FMR1, which regulates synaptic translation and can lead to fragile X syndrome when mutated, as well as the ALS-associated RBP FUS, which is involved in mRNA transport to the synapse. Interestingly, we also observed motifs mainly associated with RNA splicing (e.g., NOVA1/2 and RBFOX1/2), potentially reflecting the nuclear origin of synaptic transcripts undergoing alternative splicing before their localization (Lipscombe, 2005) or local moonlighting functions of these RBPs (Racca et al., 2010).

The differential availability of RBPs in a cell can potentially lead to different localization patterns, even when cis-acting elements in UTRs remain the same. To examine this, we extracted the expression data for the RBPs corresponding to the 3’UTR motifs detected in excitatory and inhibitory synapses from single-soma sequencing data (J. D. Perez, Dieck, et al., 2021) (Figure 3E). We found that RBP mRNA levels were highly conserved across types (∼95%), suggesting that if RBP mRNA level differences play a role in the differential localization, they are likely small, at least for the RBPs identified here. Together, this indicates that while the synaptic transcriptome is not a passive reflection of the somatic transcriptome (Figure 2), differential mRNA levels between cell types probably contribute more to the differential localization of mRNAs than changes in the 3’UTR regulatory landscape.

### Inhibitory subtype-specific synaptic transcriptome signatures

We next asked whether synaptic transcriptomes are also different between inhibitory synapse subtypes. To assess this, we crossed PV-, SST- and VIP-Cre driver lines (Figure 4A) with SypTOM mice to fluorescently label synapses in these three non-overlapping inhibitory neuron populations, which make up a large fraction of inhibitory neurons in the cortex (Tremblay et al., 2016). To study type-associated transcriptome differences, we prepared PV, SST and VIP synaptosomes from the cortex, sorted them using FASS and performed RNA sequencing. We reliably detected more than 10000 types of mRNAs per synaptosome type (Figure 4B), similar to what we observed for Camk2a and Gad2 synaptosomes (Figure 1E/F). Using PCA, we found that PV, SST and VIP synaptosomes did indeed show different transcriptomic signatures, and differed from their controls (Figures 4C and S5A). To further examine these differences, we performed differential expression analysis, comparing the inhibitory synaptosome transcriptomes to a combined control and one another (Figure 4D). We found thousands of differentially expressed mRNAs in these comparisons: ∼ 2000, 500 and 1000 mRNAs were enriched in PV, SST, VIP synaptosomes, respectively (Figure 4E). SST and VIP synaptic transcriptomes were more similar than SST/VIP and PV synaptic transcriptomes (Figure 4D bottom). We found that several established pan-inhibitory neuron markers like *Gad1, Slc32a1*, and *Gabra1* were enriched to different extents across the inhibitory synapse types (Figure 4F left); we also observed the enrichment of subtype markers like *Calb2* and *Htr3a* in VIP synaptosomes. Given the different postsynaptic targets of PV, SST and VIP neurons, we also examined the enrichment of known excitatory markers like *Camk2a*, *Arc* and *Shank2* (Figure 4F middle). As expected, PV and SST synaptosomes showed enrichment for selected excitatory markers – likely reflecting the fact that PV and SST neurons synapse with excitatory neurons – which were not enriched in VIP synaptosomes, which primarily target other inhibitory neurons (Tremblay et al., 2016). We also found several mRNAs uniquely associated with one inhibitory synapse type, as defined by enrichment in one type and negative fold change for this mRNA in both other types (Figure 4F right). Notably, the PV-enriched *Dmd* is associated with the symptoms of intellectual disability in Duchenne muscular dystrophy (Vaillend et al., 2025), the SST-enriched *Ndn* is associated with Prader-Willi syndrome (Soeda et al., 2023) and the VIP-enriched *Trpc4* is associated with autism (Jeon et al., 2025), underscoring potential synapse type-specific disease vulnerabilities.

**Figure 4:**
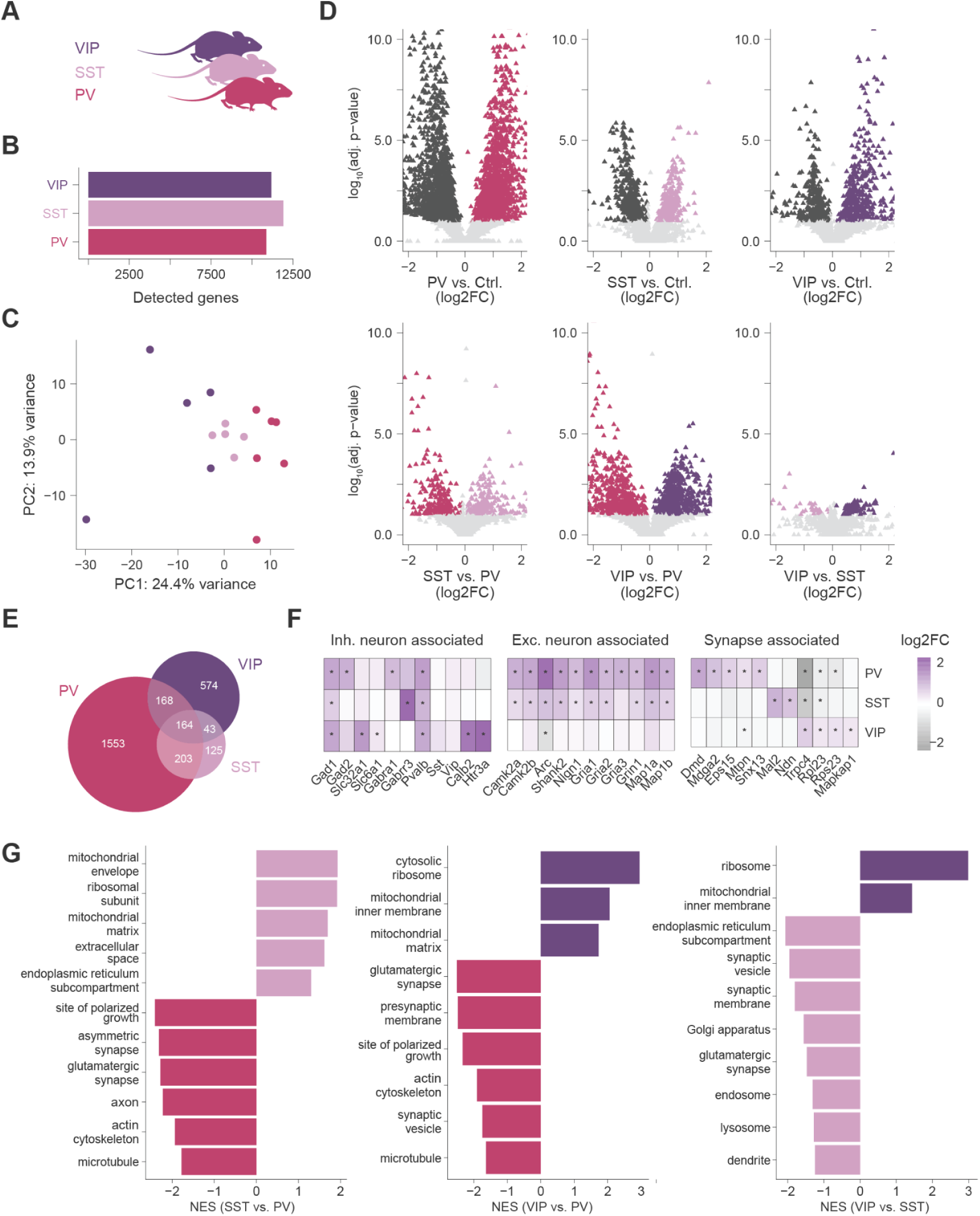
Transcriptome diversity of different inhibitory synapse types. (A) Cre-driver lines that were crossed with SypTOM mice to label synapses in PV, SST and VIP neurons (generating PV-Cre::SypTOM (PV), SST-Cre::SypTOM (SST) and VIP-Cre::SypTOM (VIP) mice). (B) Detected types of mRNAs cortical inhibitory subtypes. (C) PCA for the 1000 most variable mRNAs of PV, SST and VIP synaptosomes, showing distinct transcriptomes. (D) Volcano plot comparing PV, SST and VIP synaptosomes vs. control (top), and against one another (bottom). Significant mRNAs are colored in dark grey (control) or according to synapse type (see (A)). The x-axis is clipped at |x| = 2 and the y-axis is clipped at 10. (E) Venn Diagram of the inhibitory synapse type-enriched transcripts resulting from (D). (F) log2 fold changes of the PV, SST, VIP vs. control comparison for selected known mRNAs associated with inhibitory neuron types (left) or excitatory neuron types (middle), and identified synapse type-specific transcripts (right). (G) Selected normalized enrichment scores (NES) for terms enriched in gene set enrichment analyses conducted on ranked lists from the SST vs. PV (left), VIP vs. PV (middle) and VIP vs. SST (right) comparison.

While the transcriptome of all three inhibitory types shares lots of common features (Figure S5B), gene set enrichment analysis revealed some broader differences between them (Figure 4G). All three inhibitory synapse types showed enrichment of mRNAs associated with glutamatergic synapses with their relative enrichment (PV>SST>VIP) reflecting the increasing proportion of excitatory postsynaptic partners they interact with. mRNAs associated with metabolic processes (mitochondria or the ribosomes) also showed a gradient in enrichment (highest in VIP and lowest in SST). Finally, a key feature of PV synaptosomes was the enrichment of mRNAs associated with the cytoskeleton, while SST-enriched mRNAs were associated with the ER, Golgi apparatus and endosomes. We also assessed 3’UTR lengths and found that while PV-enriched mRNAs exhibited long 3’UTRs, uniquely enriched SST- and VIP transcripts had mean 3′UTR lengths below 1400 nucleotides (Figure S5C). Together, this shows the synaptic transcriptome can not only differentiate synaptic neurotransmitter types, but can also differentiate more subtle synaptic subtypes.

### Regional and neurotransmitter-based synaptic transcriptome signatures in the mouse brain

To what extent does the synaptic transcriptome reflect the regional diversity of synapses that are present in different brain areas? To test this, we applied our pipeline to purify nine synapse types: synapses from Camk2a- and Gad2-expressing neurons in the cortex, hippocampus, striatum, and olfactory bulb, as well as synapses from Gad2-expressing neurons in the cerebellum (Figure 5A). Using RNA sequencing, we reliably detected ∼11000 types of mRNAs in all synapse types, with 100s of mRNAs enriched in each type (Figure 5B) and a clear separation between transcriptomes of synaptosomes derived from Camk2a neurons, Gad2 neurons and controls (Figure S6A). Combining the mRNA enrichment across all comparisons (and the cortex experiment shown in Figure 1), we identified 2407 enriched mRNAs, reflecting the broad synapse diversity in these brain regions. In general, we observed that the overlap in synaptic enrichments was substantially greater within Camk2a or Gad2 synaptosomes than between Camk2a and Gad2 synaptosomes across the investigated brain regions (Figure 5C). Across brain regions, gene set enrichment analysis revealed the greatest similarity for inhibitory synapses in the cortex, hippocampus and cerebellum (Pearson’s r > 83%), whereas within-region comparisons revealed strong similarities between Camk2a and Gad2 synapses in the cortex and olfactory bulb (Pearson’s r > 75%, Figure S6B). Using partial least squares discriminant analysis (PLS-DA), we further observed that the synaptic transcriptome is shaped by both the parent cell type (Camk2a vs. Gad2) and brain region, as transcriptomes could be separated by both traits (Figure 5D/E). Using DAT-Cre::SypTOM mice, we also sequenced synaptosomes derived from dopaminergic projections to the striatum (Chuhma et al., 2023), and identified substantial overlap with Camk2a- and Gad2-enriched mRNAs and ∼50 DAT-unique mRNAs (Figures S6C-E).

**Figure 5:**
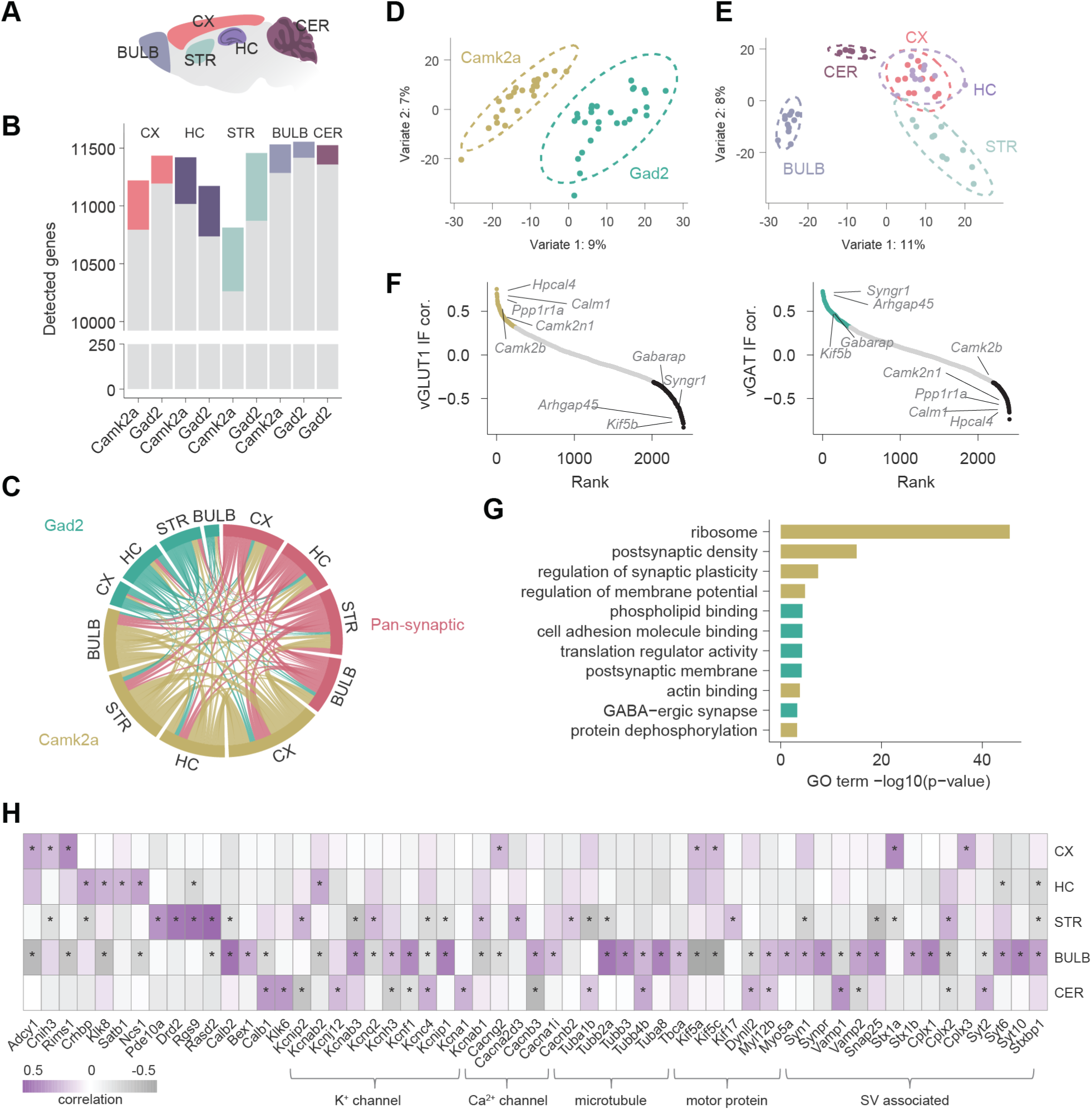
Synapse type-specific mRNA signatures across the mouse brain. (A) Schematic showing the brain regions from which synaptic transcriptomes were characterized. The following abbreviations are used - CX: cortex, HC: hippocampus, STR: striatum, BULB: Olfactory bulb, CER: cerebellum. (B) Bar plot showing the number of different mRNAs enriched (colored) and detected (grey + color) in each synapse type. 2 mio synaptosomes were sorted per synapse type sample. (C) Chord diagram showing the overlap between uniquely enriched Camk2a/Gad2-associated mRNAs and pan-synaptic mRNAs in each brain region. (D) PLD-DA of Camk2a and Gad2 synaptosomes across four brain regions (CX, HC, STR, BULB) and Gad2 synaptosomes derived from the CER. The samples were separated based on parent cell type with an overall accuracy of 92.4% (2 components, 5×50-fold CV, cendroid distance). (E) Same as (D), but separated based on brain region with an overall accuracy of 64.3%. (F) Rank plots showing the correlation of mRNAs across the mouse brain with vGLUT1/Camk2a (left) or vGAT/Gad2 (right) immunofluorescence overlap data obtained from (Oostrum et al., 2023). Significantly (p-adjust. < 0.1, Benjamini-Hochberg test) positively correlated mRNAs are colored in yellow (for vGLUT1) and cyan (for vGAT), significantly negatively correlated mRNAs are shown in black. Selected candidates are shown. (G) Barplot of p-values for selected GO terms (cellular component, biological process and molecular function) of mRNAs highly enriched with excitatory synapses (vGLUT1) or inhibitory synapses (vGAT) across the mouse brain. (H) Heatmap showing the brain region-specific association of selected synaptic mRNAs. Asterisks indicate significance (p-adjust. < 0.1, Benjamini-Hochberg adjusted). Examples for known individual brain region marker genes are shown (left), but also region-specific associations with potassium channel, calcium channel, microtubule, motor protein and synaptic vesicle mRNAs are shown (right).

Is there a set of common, correlated mRNAs associated with excitatory and inhibitory synapses across brain regions? To address this, we correlated expression levels of all 2407 synaptic mRNAs across the different synapse types with immunofluorescence data of *vGlut1/Camk2a* and *vGAT/Gad2* colocalization across the mouse brain (Oostrum et al., 2023), reflecting the extent to which inhibitory and excitatory synapses are labelled by the Camk2a and Gad2 mouse lines (Figure 5F). As expected, synaptic markers like the kinase *Camk2b* and the GABA type A receptor-associated protein *Gabarap* showed significant correlations with excitatory and inhibitory synapses, respectively. GO analysis further revealed ribosomal protein mRNAs to be a key feature of excitatory synapses across the brain (Figure 5G), allowing for potential local ribosome remodelling (Fusco et al., 2021; Shigeoka et al., 2019), or late-stage biogenesis (Fusco et al., 2025). Core mRNAs in inhibitory synapses were associated with synapse-related terms like GABAergic synapse, postsynaptic membrane and phospholipid binding (which includes mRNAs related to cytoskeletal and synaptic vesicle dynamics). Furthermore, the term translation regulatory activity was significantly enriched – several translation initiation and elongation factor mRNAs were significantly associated with inhibitory synapses, e.g., *Eef1b2*, a translation elongation factor that plays a critical role in delivering aminoacylated tRNAs to the ribosome. We further found the term cell adhesion molecule binding enriched, with several mRNAs encoding neuronal cell adhesion molecules such as *Ctntn6* and *Ank3* enriched in inhibitory synapses. Notably, microglial mRNAs like *Itgam* and *Itgb1* also contributed to the term, potentially arising from microglial processes interacting with inhibitory synapses (Z. Chen et al., 2014; Z.-P. Chen et al., 2025).

To identify distinct sets of synaptic mRNAs associated with individual brain regions, we further calculated the correlation between mRNA abundance and the brain region from which a synapse type was derived. This revealed several regional synaptic markers, including some previously described markers (e.g., the striatal marker *Rasd2)*, as well as some new synaptic markers, including sets of mRNAs related to potassium and calcium channels, microtubule and motor proteins, as well as synaptic vesicle dynamics (Figure 5H). Among the SNARE proteins, *Vamp1* was highly localized in the cerebellum and depleted in the olfactory bulb, while *Vamp2* showed the reverse pattern. *Stx1a*, in turn, was highly localized in the cortex, whereas *Stx1b* exhibited cerebellar enrichment. Together, these findings reveal that the synaptic transcriptome is shaped not only by the neurotransmitter type of the parent neuron but is also significantly influenced by regional context, indicating transcriptomic specialization that likely supports local circuit function and plasticity.

### Synaptic mRNAs are faithful markers of synapse diversity

Given the fundamental role of mRNA localization and subsequent local translation in shaping the synaptic proteome, we sought to investigate the relationship between synaptic mRNAs and proteins, using proteome data generated from the same synapse types (Oostrum et al., 2023). Many synaptic proteins are members of macromolecular complexes for which coordinated, stoichiometric synthesis of the components is desirable. As such, we asked whether mRNAs encoding members of the same complex showed coordinated synaptic localization, suggesting coordinated local translation. Indeed, we found pairwise mRNA correlations for protein complex members to be, on average, higher than for all mRNA pairs (median 63% for CORUM complexes versus 1.5% for all; Figure 6A). We next asked how strongly individual genes link their synaptic mRNA levels to protein abundance across synapse types and calculated gene-wise mRNA-protein correlations. We observed significantly higher mRNA-protein correlations than expected by chance (median 24% versus 0% for a scrambled control; Figure 6B), with 93 mRNA-protein pairs showing statistically significant (t-test, Benjamini-Hochberg adjusted p-value < 0.1) linear correlations across synapse types. These pairs, for example, comprised genes directly involved in synaptic machinery, calcium dynamics, signal transduction and neurite structure (Figure 6C), underscoring the fine-tuned coordination of mRNA-protein ratios in several genes with core synaptic functions. We also compared mRNA and protein abundances globally and observed Pearson’s correlations between 32% and 40% for all synapse types (Figure S7A), similar to mRNA-protein correlations observed in whole cells or tissue (Buccitelli & Selbach, 2020; Kaulich et al., 2025; Liu et al., 2016; Schwanhäusser et al., 2011).

**Figure 6:**
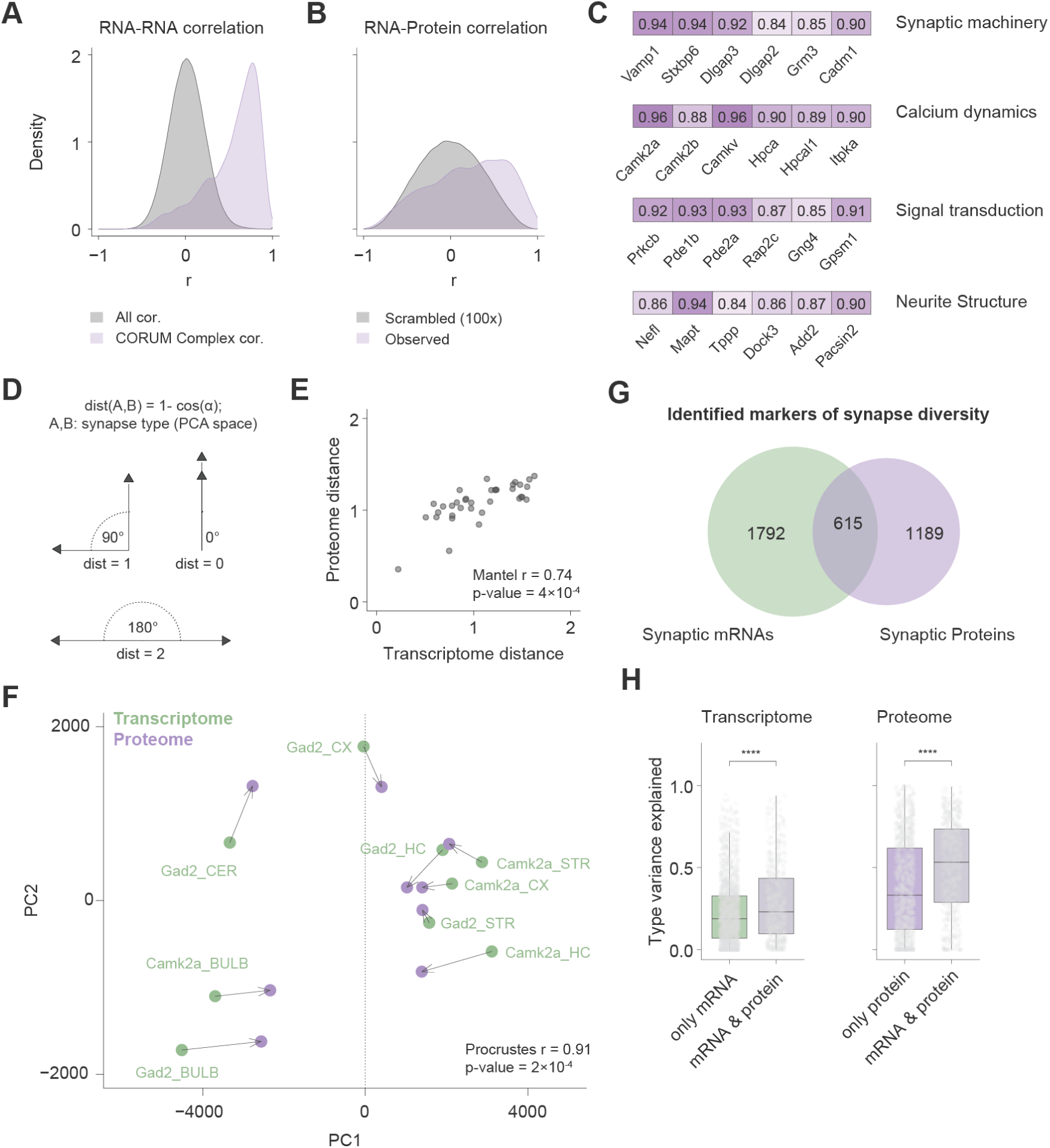
Brain-wide RNA-Protein relationship at synapses. (A) Distribution of pairwise Pearson’s correlation of all synaptically enriched mRNAs (grey) and of synaptically enriched mRNAs coding for proteins of the same CORUM complexes (purple). (B) Distribution of genewise Pearson’s correlation of mean log2-transformed mRNA and protein (iBAQ) abundances across synapse types in the mouse brain (purple) compared to a scrambled control (100x). The significant shift of the measured mRNA-protein distribution was assessed using a one-sided Wilcoxon rank-sum test (p-value < 2.2e-16). (C) Selected list of significantly correlated mRNA-protein pairs associated with the categories synaptic machinery, calcium dynamics, signal transduction and neurite structure. (D) Schematic showing examples of how the cosine distance between different samples in a PCA plot is calculated. (E) Scatter plot comparing the proteome and transcriptome distance (calculated as cosine distance) of Camk2a and Gad2 synaptosomes across the mouse brain. The first eight and 22 principal components (explains ∼90% of the variance) were used for the transcriptome and proteome, respectively. Only genes enriched either at the RNA or protein level were considered. The Mantel test for distance matrices showed significant correlation (r = 74%, p-value = 1×10⁻⁴, permutations = 10000). (F) Procrustes analysis of PCA centroids of the first 30 principal components for Camk2a and Gad2 synaptosomes across the mouse brain. Green circles show transcriptome centroids; purple circles show proteome centroids. Lines connect matched groups before and after Procrustes alignment. Only genes enriched either at the mRNA or protein level were considered. The Procrustean randomization test to assess similarity between the transcriptome and proteome PCA configurations showed significant concordance (r = 0.91, p = 2×10⁻⁴, permutations = 10000). (G) Venn diagram of mRNA and protein (Oostrum et al., 2023) markers of synapse diversity. (H) Boxplots showing the proportion of gene expression variance attributable to synapse type for three gene categories: genes reflecting synapse diversity at the mRNA level only, genes reflecting synapse diversity at the mRNA and protein level, and genes marking synapse diversity at both the mRNA and protein level. Variance partitioning was performed using linear mixed models for each gene for the transcriptome (left) and proteome (right). Genes enriched at the mRNA and protein level consistently exhibit significantly higher variance attributable to synapse type than those genes only enriched at the mRNA or protein level (Wilcoxon rank-sum test, p-value = 1.1e-06 for the transcriptome and p < 2.2e-16 for the proteome).

Is the functional diversity of synapses, as inferred from their proteome, also reflected at the level of localized mRNAs? To address this question, we performed PCA on the synaptic transcriptome and proteome (Figure S7B) and calculated transcriptome and proteome distances between synapse types across the brain (Figure 6D). For this, we used cosine distances in the low-dimensional PCA space, spanned by principal components that cumulatively explained 90% of the transcriptome and proteome diversity (Figure S7C). The correlation between the transcriptome and proteome distances was 74% (Mantel test, p = 2×10⁻⁴ for 10000 permutations), revealing substantial concordance between pairwise synaptic transcriptome and proteome distances across samples (Figure 6E). By aligning the synaptic transcriptome and proteome data in multidimensional space to find their best geometric fit, we further observed that, beyond the strong pairwise distance correlations, the overall low-dimensional structure of synapse diversity defined by the proteome is also capturedat the level of synaptic mRNAs (Procrustes correlation r = 0.91, p = 1×10⁻⁴, 10000 permutations; Figure 6F). Notably, 615 of the genes we identified as markers of synapse diversity were enriched at the mRNA and protein level (Figure 6G). More than 1000 genes were uniquely enriched at either the protein or mRNA level. This highlights that, while overall synapse diversity is conserved, the specific genes driving diversification are not necessarily the same across molecular layers. Interestingly, genes uniquely enriched in synaptic proteomes were associated with endomembrane trafficking and the proteasome, while many genes uniquely enriched in synaptic transcriptomes were associated with RNA metabolism or DNA binding (Figure S7D). To assess the extent to which the expression variance of genes across these three groups can be explained by the synapse type, we performed variance partitioning analyses of the synaptic transcriptomes and proteomes (Figure 6H). We found in both the transcriptomes and proteomes that synapse diversity marker genes enriched at both the mRNA and protein level exhibited significantly higher variance attributable to synapse type than those genes that were only enriched at one of those levels. Overall, our integrative analyses establish that, at the bulk level, synaptic mRNAs are faithful markers of synapse diversity. Beyond the significant concordance of transcriptome and proteome diversity, high pairwise mRNA correlations among protein complex members, conserved mRNA–protein ratios across synapse types, and the observation that shared mRNA and protein markers explain more variance than unique markers, all support a central role for local translation in shaping synapse diversity.

## DISCUSSION

Synapses exhibit profound functional diversity, resulting from a convergent interplay of factors such as neurotransmitter type, pre- and postsynaptic cell, electrophysiological activity, plasticity, regional identity, and interactions with non-neuronal cells (O’Rourke et al., 2012). These functional differences give rise to striking molecular synapse diversity, reflected at the level of protein abundances and even lifetimes (Bulovaite et al., 2022; Cizeron et al., 2020; Krueger-Burg, 2025; Oostrum et al., 2023; Oostrum & Schuman, 2024). While recent studies have profiled the enriched mRNAs of purified synaptosomes, these were limited to a few synapse types and excluded inhibitory synapses or lacked synapse type resolution (Hafner et al., 2019; Hobson et al., 2022; Kaulich et al., 2025; Rubio et al., 2023). With this study, we provide the first large-scale transcriptomic comparison of synapses from major excitatory and inhibitory neuron types across multiple regions of the mouse brain. We identified cell-type-specific synaptic transcriptomes from as few as two million synaptosomes, allowing us to profile transcriptomes from over 130 samples spanning 13 major synapse classes together with region- and mouse line-matched controls. In total, we detected 2407 mRNAs reflecting excitatory-inhibitory synapse diversity across the mouse brain, 53 mRNAs uniquely associated with DAT synaptosomes, and 2830 (1646 additional) mRNAs uniquely reflecting cortical inhibitory synapse subtype diversity.

Our in-depth analysis of cortical synaptic transcriptomes reveals that, beyond shared enrichments, discrete sets of mRNAs distinguish excitatory and inhibitory synapses. This is consistent with work on excitatory synapse types. For example, *ErbB4* sets off a cascade of synapse type-specific translation that promotes the formation of excitatory synapses onto PV, but not SST neurons (Bernard et al., 2022), and NMDAR-dependent local translation specifically sustains high-frequency neurotransmission of excitatory to excitatory synapses (Wong et al., 2023). Similarly, work on developing cortical projection neurons has also shown specialized growth cone transcriptomes across neuronal types (Veeraraghavan et al., 2024). In contrast, profiling single dendrites of cultured hippocampal neurons has revealed limited or difficult-to-resolve dendritic transcriptome diversity (J. D. Perez, Dieck, et al., 2021). This discrepancy likely reflects technical limitations; however, at the scale of entire dendrites, where excitatory and inhibitory inputs converge, subcellular transcriptome diversity might also be averaged out.

The unbiased profiling of mRNAs allowed us not only to identify candidates encoding proteins traditionally deemed synaptic, but also to detect mRNAs encoding proteins whose destined localization is not at the synapse. For instance, we identified several synaptically localized mRNAs at low copy numbers that encoded nuclear-localized proteins. These include *Crtc1,* the transcriptional co-activator of *CREB*, and the endosomal adaptor *Appl1*, both involved in pre- and postsynaptic synapto-nuclear signaling, respectively (Herbst & Martin, 2017; Y. Wu et al., 2020). Other candidates, such as the Mef2 MADS-box transcription factors (*Mef2a*, *Mef2c*), the SRY-related HMG-box *Sox4*, and the histone methyltransferase *Kmt5a*, have yet to be studied for their localized functions or retrograde transport.

Another common feature of local transcriptomes is the high abundance of ribosomal and mitochondrial mRNAs (J. D. Perez, Fusco, et al., 2021). In line with our findings on synaptically retained cell type diversity, a previous analysis of SynGO-annotated synaptic genes in single-soma RNA sequencing data found mRNAs encoding pre- and postsynaptic proteins to faithfully capture cellular diversity, while metabolism-related mRNAs – including ribosomal and nuclear-encoded mitochondrial mRNAs – did not (Adam et al., 2023; Koopmans et al., 2019). However, by directly profiling synaptic transcriptome diversity, rather than inferring it from somatic data, we found that these gene groups are actually strong markers of synapse diversity, particularly among inhibitory synapse subtypes.

Our analysis of inhibitory synapse subtypes further revealed that synaptic transcriptomes of SST and VIP neurons were more similar to each other than those of PV neurons – a pattern also observed at the proteome level (Oostrum et al., 2023). Interestingly, we identified an inverse relationship for the enrichment of GO terms related to mitochondrial function: depletion at the mRNA level and enrichment at the proteome level for PV compared to SST and VIP synapses. This might reflect neuron type-specific trade-offs between synaptic protein delivery and need-dependent local translation, given the high energy demands of fast-spiking PV neurons compared to other neuron types. PV synapses further exhibited the highest level of mRNA enrichment (2088 mRNAs) of all synapse types. Although technical explanations, including morphological differences between synapse types, cannot be fully ruled out, this is consistent with activity-dependent synaptic recruitment of mRNAs and ribosomes playing a critical role in elevating synaptic mRNA levels in PV neurons (Ostroff et al., 2002; G. Zhang et al., 2012). Future experiments using, for example, optogenetic stimulation of candidate neuron populations and subsequent synaptosome analysis will be needed to dissect how neural activity affects the synaptic mRNA and protein distribution of individual genes in a cell type-specific manner.

Our bioinformatic characterization of cis-acting elements in the 3’UTRs of localized mRNAs in cortical synapses confirmed previous findings showing that excitatory neurite-localized mRNAs tend to have extended 3’UTRs compared with non-localized mRNAs (An et al., 2008; Tushev et al., 2018). We found a similar extension of 3’UTRs in pan-synaptically enriched mRNAs and could not detect a significant difference in 3′UTR isoform usage between excitatory and inhibitory synapses, consistent with the idea that specific localization signals can act in a cell type-agnostic manner (Goering et al., 2023). Despite an overall comparable 3’UTR isoform diversity between excitatory and inhibitory synapses, uniquely enriched inhibitory mRNAs did not follow this trend and exhibited 3’UTRs that were even shorter than those of synaptically depleted mRNAs. We note that to fully resolve this, the non-neuronal contributions to synaptosomal transcriptomes should be assessed in the future as mRNAs localized to microglial processes generally exhibit shorter 3’UTRs than synaptic mRNAs (Vasek et al., 2023). RBP motif enrichment analysis further revealed an expansion of RBP interaction sites in excitatory versus inhibitory synaptic mRNAs. Future experiments will need to address whether this pattern reflects an expanded capacity for ribonucleoprotein granule associated mRNA transport to excitatory synapses compared to some inhibitory synapse types, and whether, for these synapse types, alternative transport mechanisms such as organelle hitchhiking are more pronounced (Bauer & Max, 2025; Harbauer & Schwarz, 2022; Y.-C. Liao et al., 2019). Moving forward, long-read sequencing will be crucial to overcome 3’UTR isoform detection limitations and resolve the full synaptic mRNA isoform diversity, including polyA tails, 5’UTRs, and coding isoforms.

Our parallel, brain-wide profiling of excitatory and inhibitory synapses across the mouse brain enabled a comprehensive assessment of transcriptomic synapse diversity, highlighting regional- and neurotransmitter-specific signatures within key gene groups relevant to synaptic transmission and trafficking. At the level of individual somata, approaches such as PatchSeq have demonstrated a clear correspondence between functional diversity and mRNA abundance (Cadwell et al., 2016). Extending this principle to the synapse and using the proteome as a readout of functional diversity, our integrated multiomic analysis demonstrates that, at the bulk level, this relationship also holds for synapse diversity. Notably, genes enriched in both modalities captured more diversity than those enriched in either the transcriptome or proteome alone. Transcriptome-proteome comparisons are often confounded by method-specific biases (Liu et al., 2016); however, the strong concordance observed in our synaptic data supports the idea that local translation is a driver of synapse diversity and establishes localized mRNAs as faithful markers of synapse diversity (Daskin et al., 2025). Despite this strong concordance at the bulk level, we expect the underlying mRNA and protein distributions to be markedly different, with mRNAs more sparsely and dynamically populating synapses in a type-specific manner. Single-synapse RNA sequencing approaches **–** though technically challenging (Hobson & Herzog, 2023) **–** especially when supplemented with in situ translation studies, will be essential for understanding how synaptic mRNAs shape the proteomes of individual synapses. Furthermore, fluorescence double-labeling of pre- and postsynapses could further reveal the role of local translation in shaping connectivity.

It is important to note that our approach depends on the purification of synaptosomes and such biochemical preparations introduce unique biases in the size and density of the selected particles, also reflected at the molecular level (Gulyássy et al., 2020). While extensive EM-based validations have established the comparability of our synaptosome preparation across synapse types (Oostrum et al., 2023), we cannot exclude the possibility that the full molecular diversity of particular synaptic populations is not captured here. We therefore propose, for future studies, to also use complementary, candidate-based approaches for individual genes (F. Zhu et al., 2018), as well as transcriptome- and proteome-wide approaches using proximity labeling. At the same time, our approach excels in throughput and compatibility with a wide range of molecular readouts from the same starting material and can be readily extended to study mRNA-protein dynamics in vivo across development, disease models, and behavioral states. We anticipate that this resource will be widely useful for studying local translation and synapse biology more broadly, and will serve as a springboard for understanding the molecular logic of gene expression across neuronal compartments.

## MATERIALS & METHODS

### Experimental model and animal details

The procedures involving animal treatment and care were conducted under the institutional guidelines which are in line with national and international laws and policies (DIRECTIVE2010/63/EU; German animal welfare law, FELASA guidelines) and approved by and reported to the local governmental supervising authorities (Regierungspräsidium Darmstadt). Specifically, euthanization was performed according to annex 2 of §2 Abs. 2 Tierschutz-Versuchstier-Verordnung.

All mice were housed in standard cages under conditions approved by the local governmental authorities, with a 12 h : 12 h light-dark cycle and food and water ad libitum. The following mouse Cre-driver lines were crossed with the SypTOM (JAX strain #: 012570, Ai34D) mouse line to fluorescently label presynaptic terminals: Camk2a-Cre (Tsien et al., 1996) (JAX strain #: 005359), Gad2-Cre (Taniguchi et al., 2011; Tsien et al., 1996) (JAX strain #: 010802), PV-Cre (Hippenmeyer et al., 2005) (JAX strain #: 008069), SST-Cre (Taniguchi et al., 2011) (JAX strain #: 013044), VIP-Cre (Taniguchi et al., 2011) (JAX strain #: 010908) and DAT-Cre (Bäckman et al., 2006) (JAX strain #: 006660). Adult mice of both sexes from the respective crosses were used for synaptosome preparations and sequencing analyses.

### Percoll density gradient-based synaptosome preparation

Synaptosome preparation was performed as described previously (Westmark et al., 2011), exactly as for the proteomic analysis of the same synapse types studied here (Oostrum et al., 2023) and for the molecular characterization of hippocampal synapses (Kaulich et al., 2025). Mice were decapitated after anesthetization with isoflurane, and the brains were immediately microdissected to isolate regions of interest (cortex, hippocampus, olfactory bulb, cerebellum, and striatum). The tissue was homogenized in a 1.0 mL glass WHEATON® Dounce Tissue Grinder on ice, using 10 strokes with a loose pestle followed by 10 with a tight pestle in gradient medium (GM, consisting of 0.25 M sucrose, 5 mM Tris-HCl, 0.1 mM EDTA, Millipore Protease inhibitor cocktail III (539134) at a 1:750 dilution, and SUPERase•In™ RNase Inhibitor (CAM2696) at 1:80). After a 10-minute centrifugation at 1,000 g and 4°C, the supernatant was applied to a Percoll density gradient (23%, 10%, and 3% Percoll in GM) and centrifuged for 5 minutes at 32500 g in a Beckman Coulter JA-25.50 rotor, using an Avanti J-26S XPI centrifuge (Beckman Coulter) set to maximum acceleration and minimum deceleration. The layers formed were designated as F0 (top), F1, F2/3, and F4 (bottom). The crude synaptosome fraction (layer F2/3) was extracted using a syringe with a blunt cannula for subsequent FASS.

### Fluorescence-Activated Synaptosome Sorting (FASS)

Synaptosomes isolated from various brain regions (cortex, hippocampus, striatum, olfactory bulb, and cerebellum) of Camk2a-Cre::SypTOM, Gad2-Cre::SypTOM, PV-Cre::SypTOM, SST-Cre::SypTOM, VIP-Cre::SypTOM and DAT-Cre::SypTOM mice were purified using FASS (Biesemann et al., 2014), closely following our previous FASS protocol using a BD FACSAriaTM Fusion Cell Sorter (SN: R656700G5029) (Kaulich et al., 2025; Oostrum et al., 2023). In brief, we loaded particles from the crude synaptosome fraction (Percoll gradient layer F2/3) and passed them through a 70 µm nozzle. Doublet particles were excluded based on SSC-H and SSC-W, and synaptosomes were purified based on either the fluorescent membrane dye (FM^TM^ 4-64 Thermofisher T13320) staining only (P2/control sample) or both membrane dye staining and Synaptophysin-tdTomato signal (P3/purified sample). Above noise levels for tdTomato were determined by comparison with synaptosomes derived from wild type mice (Figure S1A). BD FACSFlow (12756528) or PBS (Severn Biotech Ltd.,20-74-100) served as sorting buffers. Synaptosomes were collected using a tube connecting the device to a glass fiber filter (Whatman GF/F glass microfiber filter (1825-090)), which we previously rinsed with 700 µL of activated RNAsecure (ThermoFisher AM7006), diluted 1:10 in the respective sorting buffer. After collecting ten (focused Camk2a vs Gad2 cortex comparison) or two million synaptosomes (all other samples), the filter was washed with 1 mL of RNAse-free PBS buffer to collect the remaining synaptosomes and remove residual sorting buffer. The filter was subsequently stored in a 0.2 mL DNA LoBind PCR tube at −76 °C until further use.

### Library preparation and sequencing

For the preparation of sequencing libraries, we used a previously established low-input sample protocol (Kaulich et al., 2025) modified from a subcellular RNA sequencing protocol (J. Perez & Schuman, 2022). Unless otherwise stated, samples were kept on ice. First, we resuspended synaptosomes from the sorting filter by adding 7.5 µL of 1 mM DTT, 1U/µL Recombinant RNAse Inhibitor, 1X SingleShot Lysis Buffer, and 1X DNAse solution to each sample. For membrane lysis and mtDNA digestion, we incubated the samples for 20 min at 20°C, followed by 5 min at 75°C to inactivate the enzyme. We next hybridized RT primers (carrying a sample-specific index and UMI sequences) to polyA stretches in the sample RNA by adding 4.6 µL of the Pre-RT Mix (Table 1), incubating for 5 min at 65°C, and then immediately keeping the sample on ice for at least 1 min. We then carried out reverse transcription and template switching, adding the RT and TS mix (Table 1) and incubating for 10 min at 55°C, before inactivating the reverse transcriptase SuperScript IV for 10 min at 80°C. For PCR, we added the KAPA HiFi HotStart Ready Mix (1X), containing the DNA polymerase, and PCR primer (0.1 µM), bringing the samples to a final volume of 48 µL. After incubating for 3 min at 98°C for activation of the DNA polymerase, samples went through the following incubation cycles: 18 (for ten million synaptosomes) or 21 (for two million synaptosomes) cycles of incubation at 98°C for 20 s for DNA denaturation, 67°C for 10 s for annealing, and 72°C for 6 min of extension. Finally, the samples were incubated at 72°C for an additional extension phase and then immediately kept on ice. We next removed the sorting filter by transferring the sample to a centrifuge tube filter (Spin-X Centrifuge 764 Tube Filter 0.45 µM Pore CA Membrane in 2.0 ml Tube; Corning, 8162), centrifugating for 1 min at 3000 g, and then transferring the volume to 0.2 mL PCR tubes. To purify DNA and select large fragments, we used 28.8 µL (0.6X of the sample volume) AMPure XP beads (Beckman Coulter A63881) and followed the manufacturer’s instructions. Libraries with different RT primer indexes were then pooled and further indexed using the Nextera XT DNA library prep kit (Illumina) with a custom P5 primer. Paired-end sequencing was performed using the NextSeq^TM^ 2000 system (Illumina) with NextSeq^TM^ 1000/2000 P2 or P3 cartridges (100 cycles) and a custom Read 1 sequencing primer, using these read parameters: Read 1 = 26 bp, Read2 = 82 bp, Read 1 Index = 8 bp, Read 2 Index = 0. Data from two sequencing runs were combined for the brain-wide synaptosome comparison. Reaction details are shown in Figure S2. Oligonucleotide sequences and reagent details can be found in Table S1.

**Table 1:**
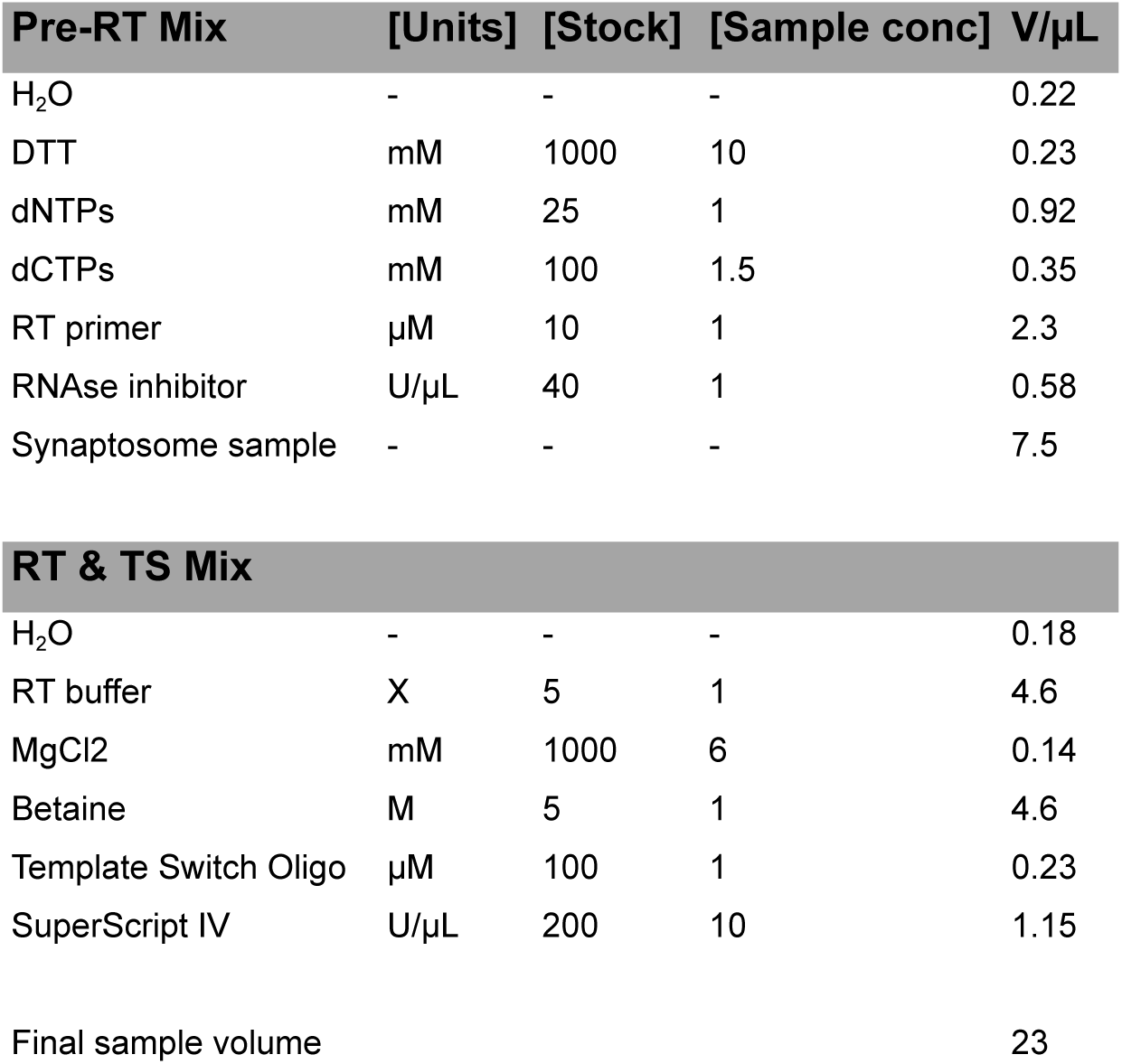
Reagent volumes and concentrations for the reverse transcriptase reaction.

| Pre-RT Mix | [Units] | [Stock] | [Sample conc] | V/μL |
| --- | --- | --- | --- | --- |
| H<sub>2</sub>O | - | - | - | 0.22 |
| DTT | mM | 1000 | 10 | 0.23 |
| dNTPs | mM | 25 | 1 | 0.92 |
| dCTPs | mM | 100 | 1.5 | 0.35 |
| RT primer | μM | 10 | 1 | 2.3 |
| RNAse inhibitor | U/μL | 40 | 1 | 0.58 |
| Synaptosome sample | - | - | - | 7.5 |

| RT & TS Mix |  |  |  |  |
| --- | --- | --- | --- | --- |
| H<sub>2</sub>O | - | - | - | 0.18 |
| RT buffer | X | 5 | 1 | 4.6 |
| MgCl<sub>2</sub> | mM | 1000 | 6 | 0.14 |
| Betaine | M | 5 | 1 | 4.6 |
| Template Switch Oligo | μM | 100 | 1 | 0.23 |
| SuperScript IV | U/μL | 200 | 10 | 1.15 |

|  |  |  |  |  |  |
| --- | --- | --- | --- | --- | --- |
| Final sample volume |  |  |  |  | 23 |

### Preprocessing methods for RNA-Seq of synaptosome datasets

For preprocessing, we used a previously published custom bash pipeline for barcoded bulk synaptosome RNA sequencing (Kaulich et al., 2025; Spanò et al., 2025). In short, raw BCL files were demultiplexed with bcl2fastq, generating paired-end FASTQ files. Paired-end reads were aligned to the Mus musculus mm39 reference genome and annotation from the UCSC Genome Browser using STAR (v2.7.11b) (Dobin et al., 2012) in STARsolo mode (--soloType CB_UMI_Simple). Cell barcodes (one mismatch allowed against a predefined whitelist, --soloCBmatchWLtype 1MM) and UMIs were parsed from read 1 and UMIs were deduplicated with a one-mismatch scheme across all genes (--soloUMIdedup 1MM_All). Gene expression quantification was performed with --soloFeatures GeneFull_Ex50pAS. Resulting BAM files, UMI count matrices and alignment logs were saved for subsequent analyses.

### Analysis of synaptic transcriptomes

After generating RNA count matrices, we conducted further analysis in R 4.4.0. We focused our analysis on protein-coding RNAs (based on the Mus musculus mm39 reference genome, the NCBI RefSeq assembly: GCF_000001635.27) to characterize potential templates for local protein synthesis. Transcripts were filtered (on average, at least one count per sample) to remove potential noise. Counts originating from mitochondrially encoded genes and high-confidence microglial contaminants (see single-soma RNA sequencing data analysis) were further removed to focus on localized transcripts in neurons whose transcription occurs in the cell nucleus. We then corrected for Nextera PCR batch effects using COMBATseq (sva package v3.52.0) (Y. Zhang et al., 2020). For downstream analyses, we used DESeq2 1.44.0 (Love et al., 2014) to construct a DESeqDataSet object using the factor “Type” – encoding the different sorted synapse types and control samples within and across brain regions – as the design formula (design = ∼ Type).

For dimensionality reduction of synaptic transcriptomes, counts were normalized blind to sample group (blind=TRUE) using the variance-stabilizing transformation (vst) in DESeq2. Normalized counts were then used as input for different dimensionality reduction methods. We used the prcomp function from the R “stats” package to perform PCA. Outliers were removed based on two-dimensional PCA plots. To perform PLS-DA on synaptic transcriptomes across the mouse brain, we used the mixOmics package (Rohart et al., 2017), separating transcriptomes based on the factors “region” and “parent cell type” as class variables. The mixOmics perf function was used to evaluate PLS-DA model performance using 5-fold cross-validation (50 repeats).

We performed differential expression analysis in DESeq2, using a negative binomial generalized linear model with the default Wald test and local dispersion fit for the Camk2a vs. Gad2 synaptosome comparison, and a parametric dispersion fit for the PV, SST, VIP synaptosome comparisons. Log2 fold changes were shrunk using the apeglm method (A. Zhu et al., 2018). DESeq2 p-values were adjusted using independent hypothesis weighting (IHW package v1.32.0) at a significance level of 0.1 (Ignatiadis et al., 2016). As is standard practice, we modeled the weight of each test using the mean of normalized counts as a covariate.

We defined synaptically enriched mRNAs as follows. Cell type-specific synaptic transcriptomes were compared to their respective controls, and mRNAs were deemed enriched if log2 fold change > 0.1 and the adjusted p-value < 0.1 (and depleted for log2 fold change < −0.1 and adjusted p-value < 0.1). Since we did not observe mouse line-specific differences, control samples were pooled within brain regions. Cell type-specific synaptic transcriptomes were also directly compared to one another, and mRNAs significantly enriched in such comparisons were also defined as synaptic, as long as they didn’t show a negative log2 fold change compared to their respective controls. In the case of synaptic transcriptomes derived from DAT-Cre::SypTOM mice, we only compared them to their controls because we previously found them to be less pure than Camk2a and Gad2 synaptosomes (Oostrum et al., 2023). For cortex-enriched synaptic mRNAs (Camk2a and Gad2 neurons), enrichments from two experiments (cortex only and the brain region comparison) were pooled.

### Functional enrichment analysis

GO enrichment analysis was performed using the clusterProfiler package (v4.12.6) (T. Wu et al., 2021) with the Mus musculus annotation database org.Mm.eg.db. Lists of enriched mRNAs were tested against a background comprising all detected mRNAs after count filtering (mean count > 1 across samples) or the common default background was used (for DAT enrichment comparison). P-values were calculated using a Fisher’s exact test and were subjected to Benjamini-Hochberg multiple-testing correction.

GSEA was performed using the clusterProfiler::gseGO function. Ranked lists of mRNAs were used as input for the analysis **–** either based on correlation values or differential expression analysis results (−log10(adjusted p-value) x sign(log2 fold change)). For each GO term, p-values were calculated by comparing the enrichment scores to scores from 1000 random permutations. Multiple-testing correction was done using Benjamini-Hochberg and significant normalized enrichment scores were reported.

### Single-soma RNA sequencing data analysis

To obtain mRNA abundances, define cell type markers, and assess to what extent synaptically enriched mRNAs reflected parent cell type identity, we analyzed a previously published single-soma RNA sequencing data set of the mouse isocortex and hippocampal formation (Yao et al., 2021). Count matrices were log-normalized and differential expression analysis between annotated classes was performed using Scanpy’s rank_genes_groups function (v1.11.5) in Python 3.14.0. Cells were grouped by the annotated class_label field, and pairwise comparisons were computed for each class against all remaining cells using the Wilcoxon rank-sum test. For every gene and class, Scanpy returned the test statistics, nominal p-values and Benjamini-Hochberg adjusted p-values. Only genes with an adjusted p-value < 0.05 were retained. To generate a unified marker table, for each class we extracted (i) the gene rank within the class-specific DE list, (ii) the minimum rank of the same gene across all other classes, and (iii) the resulting rank-separation score (Δrank). Library- and log1p-normalized mean expression values were computed directly from the AnnData object by averaging the expression of each gene within the focal class and across all other classes. To quantify class specificity, we additionally calculated τ (tau), a normalized specificity index (0-1) based on relative expression across all classes, where 1 indicates expression restricted to a single class. Microglial marker mRNAs were defined within the “Micro-PVM” group as those with Δrank > 200 and τ > 0.9. To compare mRNA preferences between somata and synapses, we computed the gene-wise differences of z-scores between the cortical synaptic expression values in this study (Camk2a, Gad2) and the mean expression values for Glutamatergic and GABAergic single somata (both log1p-normalized), respectively.

To quantify the expression of synaptosome-related gene sets at the single-cell level, we computed module scores for each synaptosome category using scanpy.tl.score_genes, restricting scoring to categories with at least three genes. The resulting per-cell module scores were compared across class_label groups. Group-wise differences were assessed using non-parametric statistics: Wilcoxon signed-rank tests were used to evaluate whether module scores within individual neuronal classes were significantly greater than zero, and Dunn’s post-hoc tests (Benjamini-Hochberg corrected) were applied to assess pairwise differences across all classes. Violin plots were generated to visualize score distributions and to annotate significant pairwise contrasts. This procedure enabled the identification of neuronal classes exhibiting selective enrichment of synaptically enriched mRNAs.

### 3’UTR and RBP motif enrichment pipeline

De novo 3′UTR isoform identification was performed using the PASSFinder pipeline (Tushev et al., 2018) based on genome-aligned reads (BAM files). To enable accurate deduplication in unannotated regions, the RefSeq mm39 gene annotation was extended 15 kb downstream of the stop codon. Reads were assigned to these extended gene coordinates with featureCounts (Subread package v2.0.8) (Y. Liao et al., 2014). BAM files were then collapsed using UMI-tools v1.1.5 (Smith et al., 2017) based on UMI, gene ID, and sample barcode, followed by quality filtering (samtools view -q 255 (samtools v1.21)). To meet PASSFinder input requirements, reads containing ≥10-nt terminal poly(A) stretches were identified based on CIGAR strings and retained. Collapsed filtered BAM files served as input to PASSFinder. A genome-wide poly(A) mask was generated using bowtie v1.3.1 (Langmead et al., 2009), and PASS detection, clustering, and annotation were performed using a modified RefSeq mm39 BED file according to PASSFinder guidelines. Only PASSFinder-validated 3′UTR isoforms were retained.

For downstream analyses, we focused on tandem 3′UTRs (96% of detected isoforms), as only 4% showed alternative last exons. First, the number of 3’UTR isoforms per gene was calculated. Because the library design produced strong 3′ end bias and short reads (82 nt), isoforms had to be sufficiently separated to ensure unambiguous assignment. Therefore, only isoforms spaced ≥170 nt apart (∼ insert size − read length) were retained. If two isoforms were spaced < 170nt, the upstream (shorter) isoform was removed from the dataset. Further strand-overlapping isoforms were removed.

To eliminate internal priming artifacts, isoforms were filtered based on the ratio of external poly(A) reads (soft-clipped ≥10 at read ends) to internal poly(A) reads. Isoforms with a ratio <10, or genes with strong internal priming within the CDS (3′UTR/CDS ratio <2.5), were excluded. Isoforms >15 kb were also removed as potential annotation artifacts.

For isoform-specific quantification, each 3′UTR was partitioned into unique sequence fragments, and the quantification window was limited to 550 nt upstream of the poly(A) site if the distance to the next upstream polyA site was > 550. Collapsed BAM files were intersected with these unique regions using bedtools intersect (-f 1.0) (bedtools v2.31.1). From the resulting count matrix, we computed gene-wise weighted average 3′UTR lengths (for all genes with >1 isoform; otherwise, single isoform lengths were used), identified the dominant isoform per gene, and performed differential isoform usage analysis using DRIMSeq v1.32.0 following the recommended analysis steps (Nowicka & Robinson, 2016). Only isoforms detected in at least 4 replicates with 5 counts each were included in the analysis.

To assess potential RBPs interacting with synaptically enriched transcripts, we performed RBP motif enrichment using AME v5.5.4 (McLeay & Bailey, 2010) from MEME Suite (Bailey et al., 2009). Synaptically enriched mRNAs were represented by their most abundant 3′UTR isoform, while control mRNAs (all mRNAs that were detected but not synapse-enriched) were represented by their longest isoform. A background model was generated with fasta-get-markov using control 3′UTR sequences, which were also supplied to AME as control input. Motifs with an AME E-value <10 (default) were considered significantly enriched. The position weight matrix collection consisted of merged and de-duplicated PWMs from ATtRACT (Giudice et al., 2016) and RCRUNCH (Katsantoni et al., 2023), yielding 1,231 motifs representing 246 RBPs.

### Genomic distribution of reads

To quantify the genomic distribution of mapped reads, we first generated a BED annotation file following the guidelines provided by PASSFinder for use with the PASSAnnotate.pl module. Using bedtools intersect, this BED6 annotation file was intersected with BAM files containing uniquely mapped and deduplicated reads. Following the intersection, reads assigned to multiple overlapping genomic features were collapsed based on their read ID and assigned to the most downstream feature. Read counts were then aggregated per genomic feature. In a final step, we generated a gene label-to-gene biotype mapping table from the mm39 refLink file to assign a biotype to each annotated gene. The resulting annotated read-count table was used to visualize the genomic distribution of reads (Figure S4).

Read coverage tracks were generated by extracting reads from specified genomic regions using samtools view -buh on collapsed BAM files filtered for uniquely mapped reads. The extracted reads were converted into bedGraph format using bedtools genomecov with the -split option to accurately account for spliced alignments. The resulting bedGraph files were subsequently converted to bigWig format using the UCSC bedGraphToBigWig utility for visualization in the UCSC Genome Browser.

### Correlation analysis to identify synaptic excitatory and inhibitory mRNA markers across brain regions

To identify core mRNAs associated with excitatory and inhibitory synapses across the mouse brain, we used immunofluorescence data (Oostrum et al., 2023), estimating the overlap the Synaptophysin-tdTomato signal in Camk2a-Cre::SypTOM and Gad2:SypTOM mice across the investigated brain regions with vGLUT1 and vGAT immunofluorescence, respectively (Table S4). For each mRNA (enriched in at least one synapse type), we computed gene-trait Pearson’s correlation between its abundance and the proportion of vGLUT1- or vGAT-labeled synapses across synapse types in the mouse brain. P-values for each correlation were obtained using a Student’s t approximation. Correlations with Benjamini-Hochberg-adjusted p-values < 0.1 were considered significant. To identify mRNAs associated with specific brain regions, we applied the same procedure, using a binary region indicator taking values 1 for synapse types originating from the region of interest and 0 otherwise. All results are shown in Table S4.

### Analysis of synaptic mRNA-protein relationships

To assess coordinated mRNA transport, we queried the CORUM complex database (Tsitsiridis et al., 2022), filtered for the organism “Mouse” and computed pairwise Pearson’s correlations across synapse types for mRNAs encoding proteins that are part of the same protein complex. Only mRNAs enriched in at least one synapse type were included in the analysis. In this and all following analyses vst-transformed RNA counts were log-normalized for better comparability with the proteome data.

To assess protein-mRNA relationships in synaptosomes generated from the same synapse types, we used our previously published DIA LC-MS/MS dataset (PXD039946) (Oostrum et al., 2023). Using the original Spectronaut (v16) file, we re-exported protein quantifications based on iBAQ values directly from Spectronaut to enable comparison between protein abundance and the corresponding RNA profiles of matched synapse types. No additional downstream statistical processing (e.g., MSstats) was applied for the present analyses. Protein gene name conversions were done using mapped IDs from UniProt.

Transcriptome-proteome-wide and gene-wise mRNA-protein correlations (Table S4) were computed using Pearson’s correlation. For the gene-wise comparison, significance was tested as described for the correlational analysis to identify excitatory/inhibitory or regional mRNA markers. The distribution of gene-wise mRNA–protein correlations was compared to a scrambled (100x) control distribution, and significance was assessed using a one-sided Wilcoxon rank-sum test.

PCA was conducted as described before for the log-normalized synaptic transcriptomes and proteomes. Transcriptomic and proteomic distances between synapse types were quantified by computing pairwise cosine distances (using principal components cumulatively explaining 90% of the variance; n_RNA_=8, n_protein_=22) and then aggregating these to distance matrices of average distances between synapse types. Cosine distance was defined as 1−cos(α), where α is the angle between sample vectors in PCA space. To test the similarity of distance matrices, we used a Mantel test (vegan package v2.7-2) with 10000 permutations to assess statistical significance. To assess overall geometric agreement between the synaptic transcriptome and proteome, PCA coordinates for individual synaptosome samples were averaged within each synapse type to obtain type-specific centroids in PCA space. Procrustes analysis was then performed on these centroids (vegan::procrustes), and significance was tested using a Procrustes randomization test with 10000 permutations (vegan::protest).

Variance partitioning (variancePartition package v1.34.0) (Hoffman & Schadt, 2016)was performed using synapse type as the explanatory factor. We focused the analysis on genes enriched at the transcriptome level, proteome level, or both in at least one synapse type, and quantified the fraction of expression variance in mRNA and protein explained by synapse type. Differences in the distributions of explained variance between gene groups were assessed using the Wilcoxon rank-sum test. All mRNA-protein comparisons were conducted using only genes enriched either at the mRNA or protein level in at least one synapse type.

### Additional info on data visualization

Basic visualization of synaptosome data was conducted in R, using ggplot2 (v4.0.0), ggrepel (v0.9.5), ggpubr (v0.6.1) and patchwork (v1.3.2). Circular chord diagrams were generated with the circlize package (v0.4.16) (Gu et al., 2014). Heatmaps were visualized using pheatmap (v1.0.13). Venn diagrams were generated with the VennDiagram package (v1.7.3). Flow cytometry data were visualized using BD’s FlowJo (v10). Read coverage tracks were inspected with the UCSC Genome Browser. Data visualization in Python was done using matplotlib (v3.10.7) and seaborn (v0.13.2). Final figures were assembled and edited for layout in Adobe Illustrator (v27.1.1).

## Supporting information

Table S2: Differential expression results for all comparisons

Table S1: Oligonucleotide sequences and reagent materials for the RNAseq protocol

Table S4: Camk2a and Gad2 Immunofluorescence values and correlation analysis results

Table S3: Single-soma RNA sequencing analysis results

## RESOURCE AVAILABILITY

The RNA sequencing data will be available via the SRA repository of NCBI upon publication. The processed data can also be accessed and visualized using our laboratory’s public interface (https://syndive.org/).

The previously established pipeline used for bioinformatic preprocessing of our RNA sequencing data is available here: https://gitlab.mpcdf.mpg.de/mpibr/schu/synaptosort. De novo 3′UTR isoform identification was performed using The PASSFinder pipeline: https://gitlab.mpcdf.mpg.de/mpibr/schu/PASSFinder. These pipelines, along with custom scripts for synaptosome RNA sequencing differential expression analysis, single-soma RNA sequencing analysis, 3′UTR isoform analysis, and RBP motif enrichment, as well as count tables and additional sample details, will be available via the Edmond repository (https://edmond.mpg.de/) upon publication (https://doi.org/10.17617/3.VMEGGS).

## ACKNOWLEDGEMENTS

We thank Dr. Elena Ciirdaeva and the MPIRBR animal facility for technical support and assistance. We thank Dr. Ashley Bourke and the rest of the Schuman lab for discussions and Julia Kuhl for graphical support. M.J. acknowledges funding from the International Max Planck Research School for Neural Circuits. E.K. is supported by EMBO (Postdoctoral Fellowship 1104 EMBO ALTF 148-2023). M.v.O. was funded by the National Science Foundation (SNSF) (P2EZP3_191820 and P400PB_199288) and the Novartis Foundation for medical-biological research (22B079). E.M.S. acknowledges laboratory funding by the Max Planck Society and the European Union (ERC, DiverseSynapse, 101054512). Neither the EU nor the granting authority can be held responsible for them.

## AUTHOR CONTRIBUTIONS

M.J. co-wrote the manuscript and performed and analyzed all experiments (with support noted below). J.M. performed bioinformatic preprocessing, de novo 3’UTR detection, and RBP enrichment analysis. J.P. conducted proof-of-principle experiments, contributed to the bioinformatic preprocessing, and supported the establishment of the RNA sequencing protocol. M.v.O. and E.K. performed FASS. S.t.D. performed synaptosome preparations and coordinated the mouse lines. N.F. performed synaptosome preparations. G.T. contributed to bioinformatic preprocessing and the single-soma RNA sequencing analysis. M.K. performed synaptosome preparations. E.M.S. supervised the project and co-wrote the manuscript.

## SUPPLEMENTARY FIGURES

**Figure S1:**
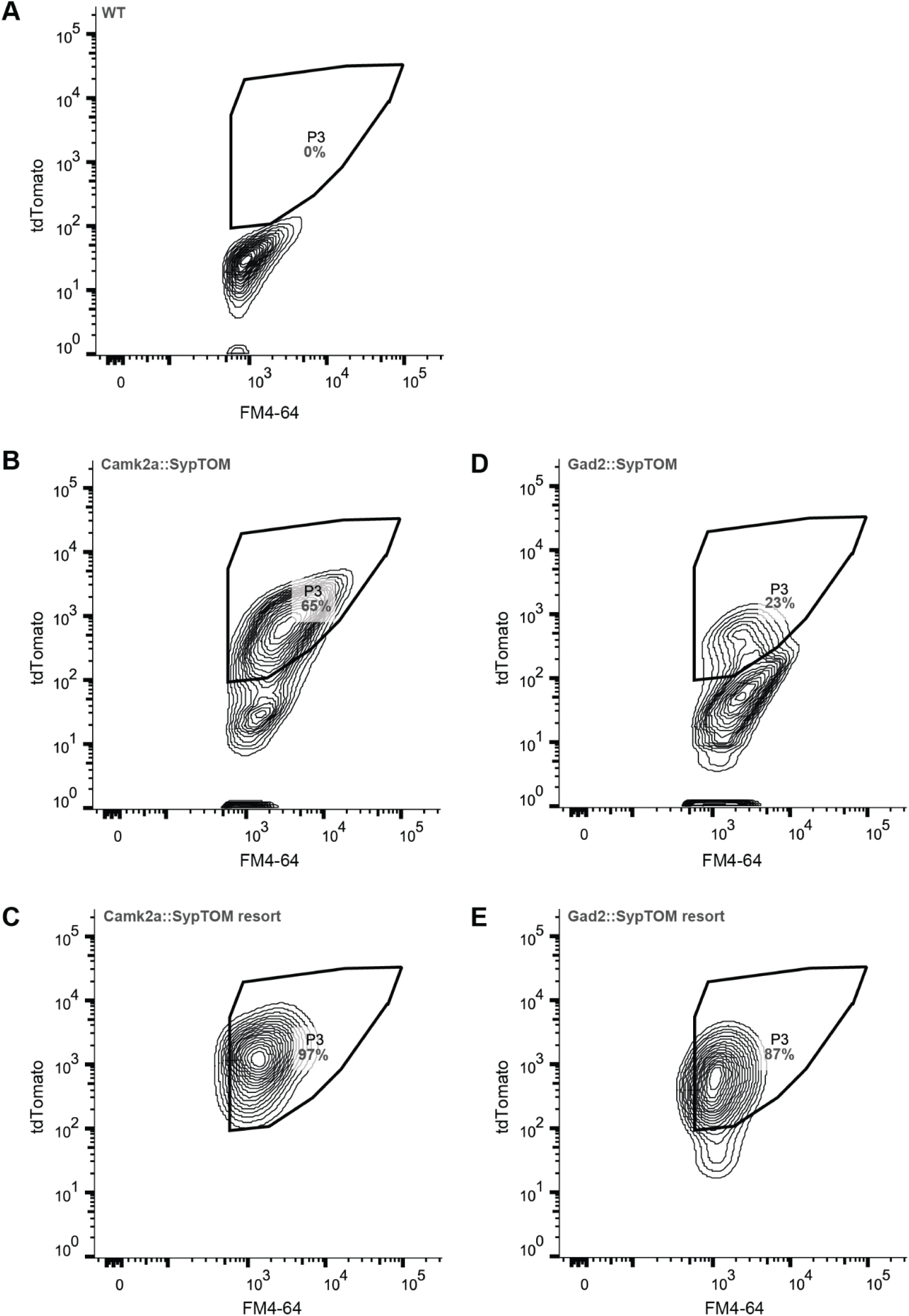
Representative FASS contour plots illustrating the gating strategy, related to Figure 1. (A) Representative contour plot (tdTomato fluorescence as a function of membrane dye FM4-64 fluorescence) for synaptosomes derived from a wild type mouse cortex. (B) same as (A), but for synaptosomes from a Camk2a-Cre::SypTOM mouse cortex. (C) Resort of sorted synaptosomes from (B). (D,E) same as (B,C), but for a Gad2-Cre::SypTOM mouse cortex.

**Figure S2:**
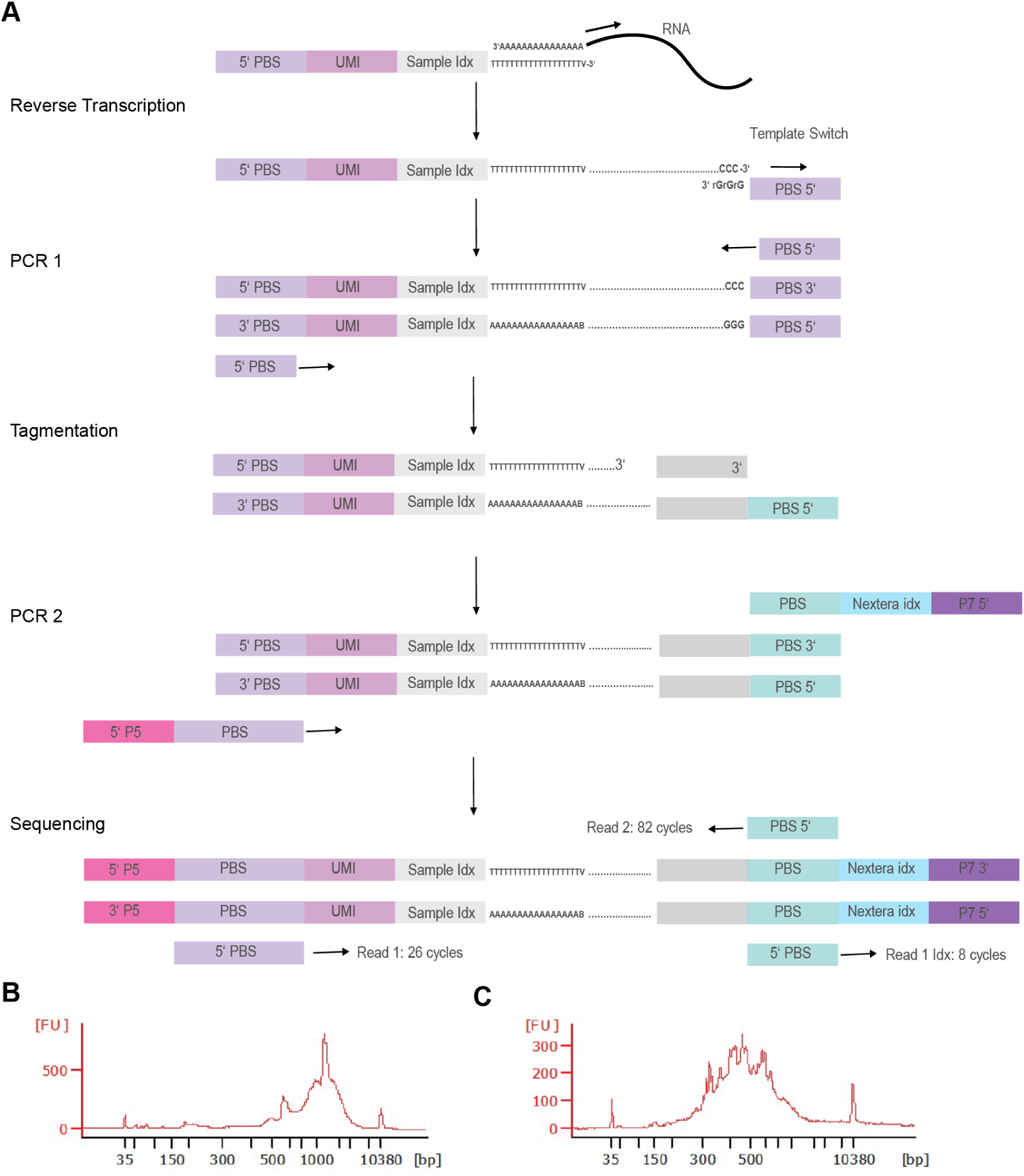
Overview of the RNA sequencing workflow, related to Figure 1. (A) Reaction scheme for the RNA sequencing library preparation and paired-end Illumina sequencing. (B) Agilent Bioanalyzer electropherogram of the full-length cDNA distribution of an exemplary library after PCR 1. (C) Agilent Bioanalyzer electropherogram of the cDNA distribution of an exemplary library after tagmentation and PCR 2.

**Figure S3:**
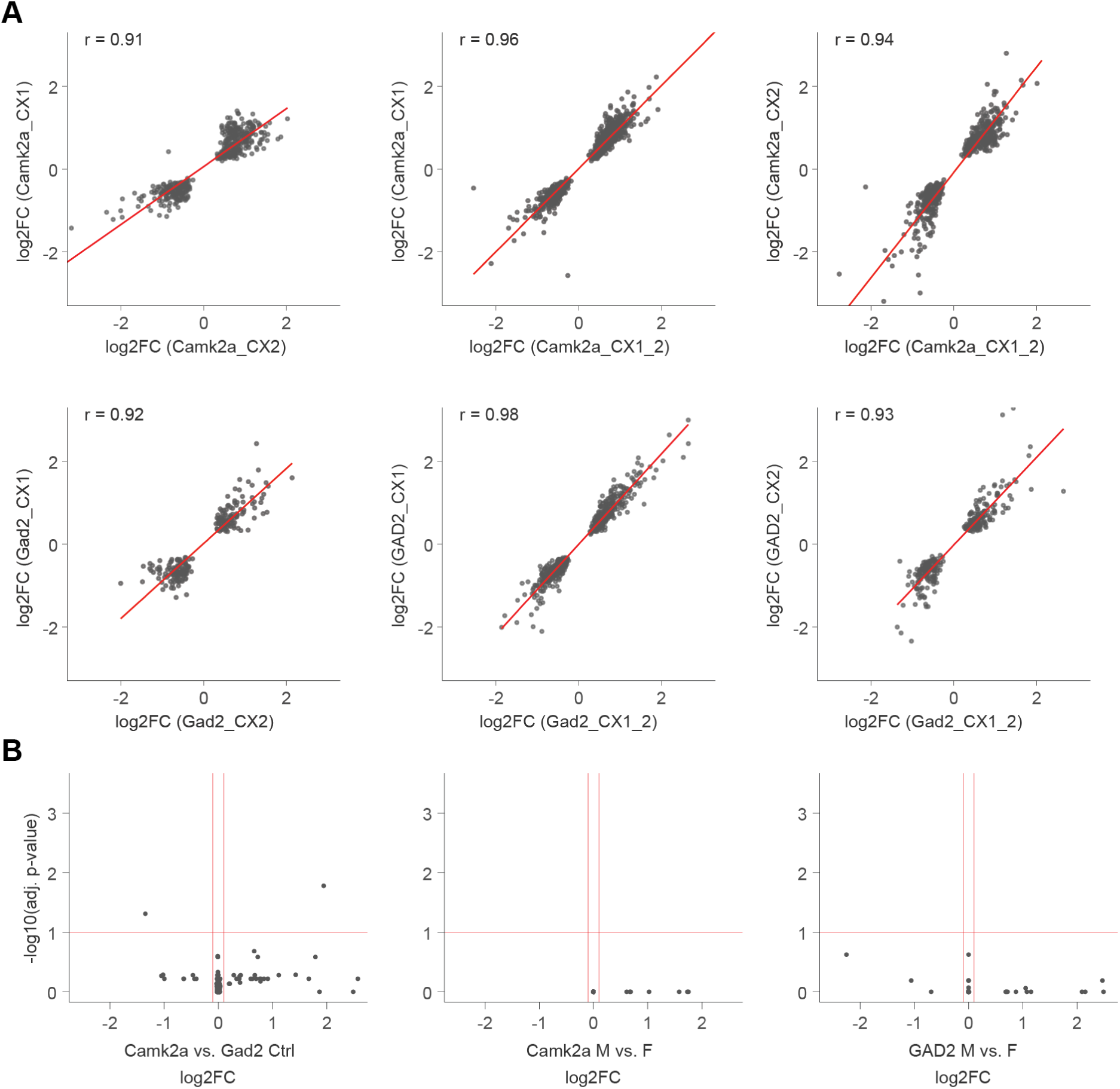
Unbiased and reproducible differential expression analysis for synapse transcriptome discovery, related to Figure 1. (A) Pearson’s correlations of significant log2 fold changes for cortex experiment 1 (CX1: 10 mio synaptosomes each, 6-8 replicates), cortex experiment 2 (CX2: 2 mio synaptosomes each, 4-6 replicates within a larger experiment comparing different brain regions) and both experiments computationally pooled (CX1_2). Correlations greater than 0.93 indicate high reproducibility of the differential expression results across litters, animals, and sequencing runs. (B) No significant transcriptome differences were observed between control samples of Camk2a and Gad2 crude synaptosome samples, underscoring the comparability of the Camk2a and Gad2 mouse lines.

**Figure S4.**
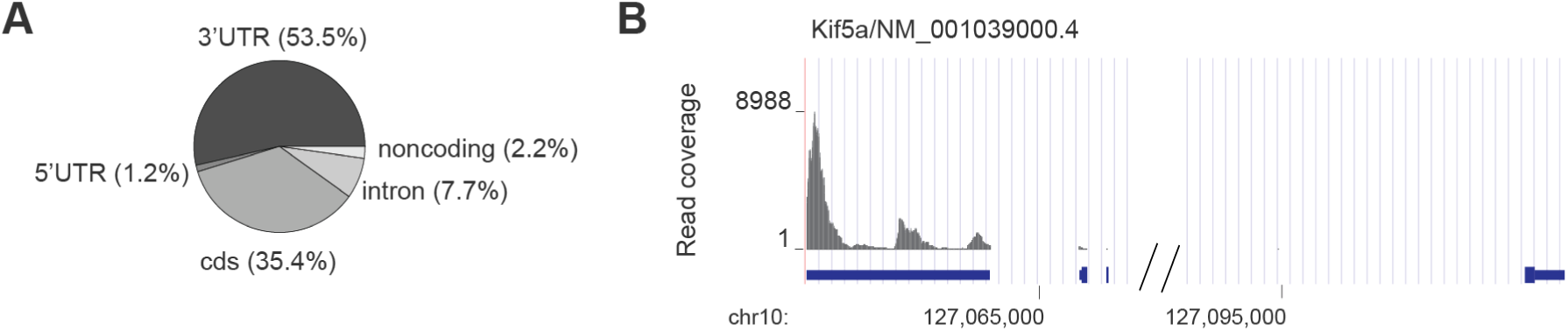
Read coverage distribution, related to Figure 3. (A) Genomic distribution (mm39) of reads, based on RNA sequencing of synaptosomes in the mouse cortex. (B) Read coverage plot for Kif5a (negatively-stranded transcript). Exon position is indicated in blue. Most reads are distributed in the 3’UTR region (three peaks, indicative of three 3’UTR isoforms).

**Figure S5:**
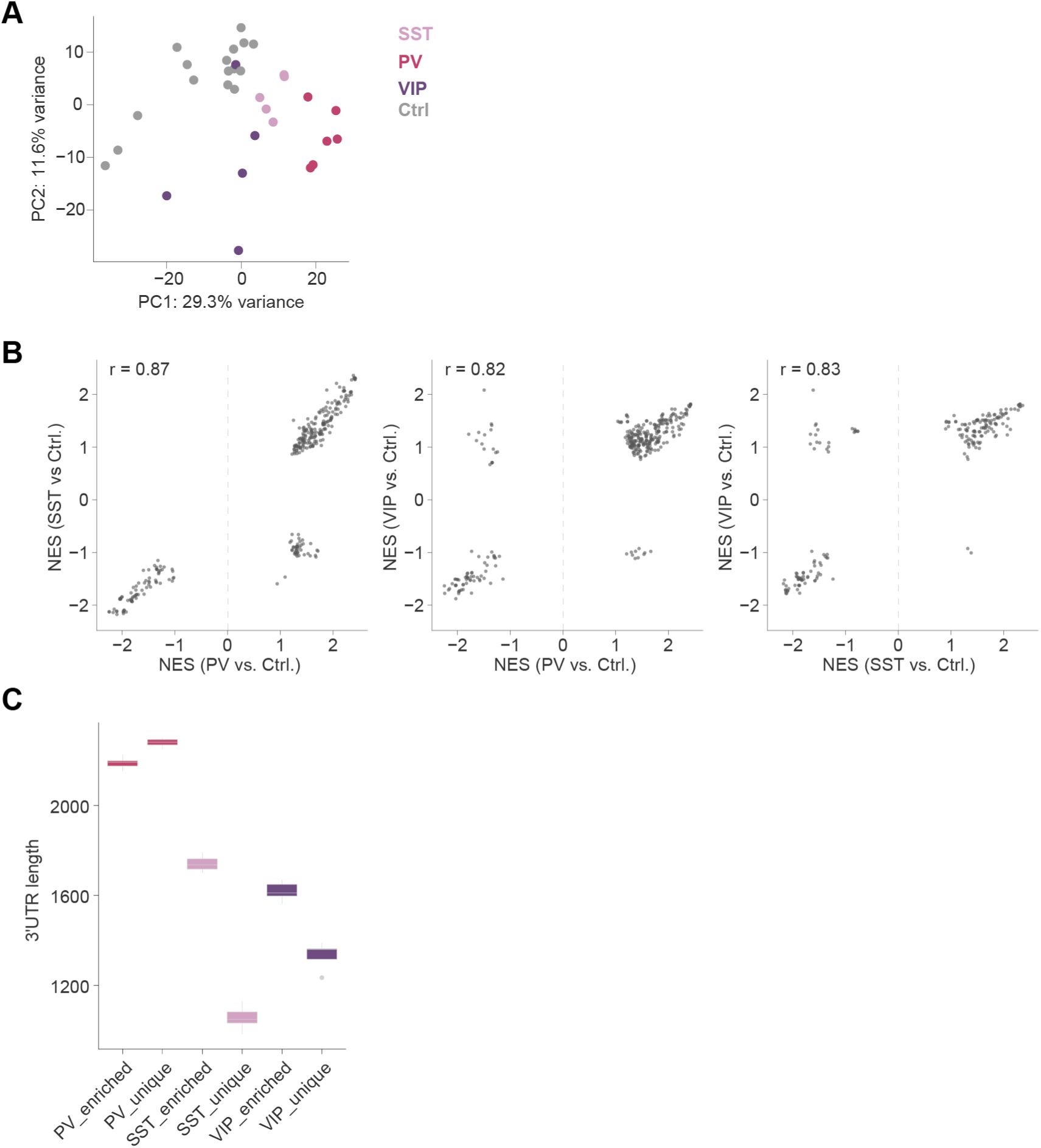
Transcriptome differences of synaptosomes derived from cortical inhibitory interneurons, related to Figure 4. (A) PCA for the 1000 most variable mRNAs of PV, SST, VIP and control synaptosomes, showing distinct transcriptomes for all particle types. (B) Results of gene set enrichment analyses (biological process, term size between 100 and 600) conducted on ranked lists from the PV vs. control, SST vs. control and VIP vs. control comparisons. Scatter plots of normalized enrichment scores (NES) are shown, comparing PV and SST enrichments (Pearson’s correlation r = 87%), PV and VIP enrichments (Pearson’s correlation r = 82%) and VIP vs SST enrichments (Pearson’s correlation r = 87%). (C) Boxplot showing the average 3’UTR length for PV, SST and VIP synaptosomes, weighted by expression of the detected 3’UTR isoforms. All categories are significantly different (one-way ANOVA p-value < 2×10⁻6; Tukey’s HSD p-value < 7×10⁻³ for all pairwise comparisons).

**Figure S6:**
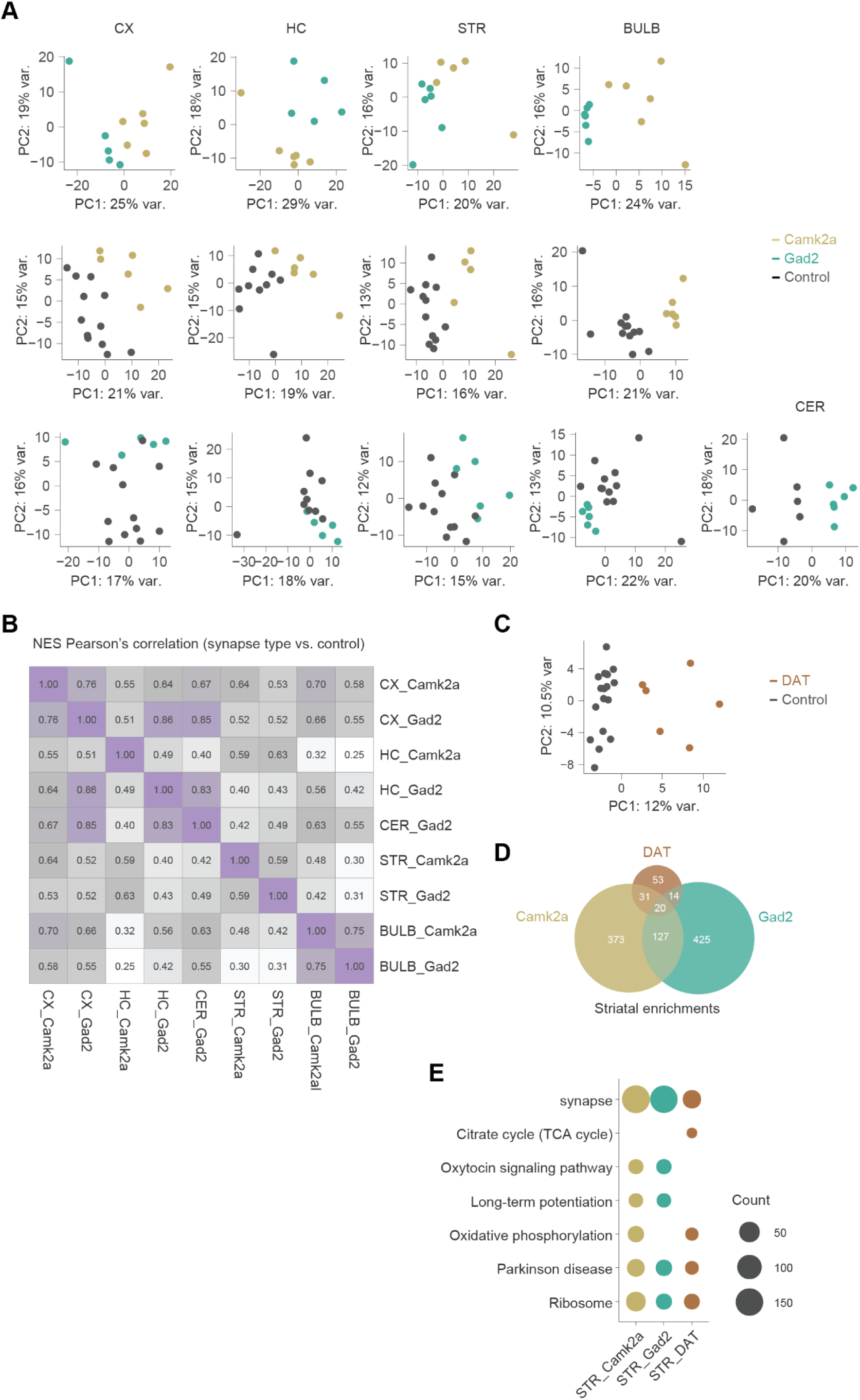
Synaptic transcriptomes across the mouse brain, related to Figure 5. (A) PCA (top 1000 mRNAs) of the nine studied synapse types comprising excitatory and inhibitory synapses across the cortex, hippocampus, striatum, olfactory bulb and cerebellum reveals clear separation between Camk2a- and Gad2 synaptosomes across the brain as well as separation from region-specific controls. (B) Gene set enrichment analysis (biological process, term size between 100 and 1000) for all synapse types. The heatmap shows the Pearson’s correlation of normalized enrichment scores (NES) calculated based on enrichment compared to each region-specific control. (C) PCA (top 1000 mRNAs) of DAT-Cre::SypTOM synaptosomes versus Ctrl. (D) Venn diagram of mRNA enrichments for Camk2a-, Gad2- and DAT synaptosomes from the striatum. (E) GO analysis (KEGG pathways and cellular component) for all mRNAs enriched in striatal synapses. The dots indicate the number of mRNAs belonging to a significant (p < 0.05) term.

**Figure S7:**
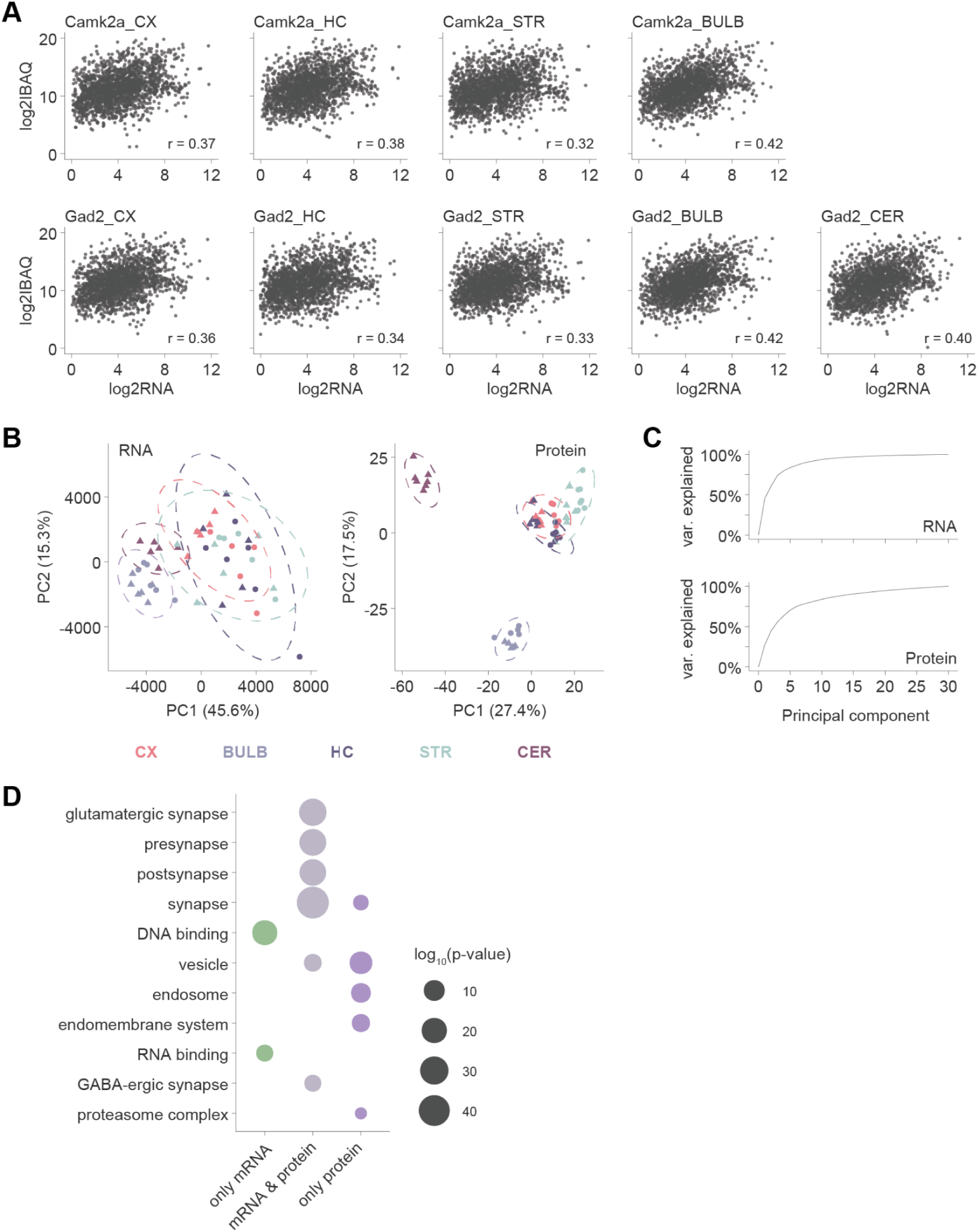
Synapse diversity at the mRNA and protein level, related to Figure 6. (A) Scatter plot and Pearson’s correlation of mean log2-transformed mRNA and protein (iBAQ) abundances of Camk2a- and Gad2-purified synaptosomes from the cortex, hippocampus, striatum and cerebellum. (B) PCA plots based on the transcriptome (left) and proteome (right) data for all nine synapses types shown in (A). (C) Cumulative variance explained as a function of the principal components for the transcriptome (top) and proteome (bottom). (D) Dotplots showing selected GO terms (molecular function and cellular component) for synapse diversity markers only enriched at the mRNA level, only enriched at the protein level or enriched at both levels.

